# No consistent effects of humans on animal genetic diversity worldwide

**DOI:** 10.1101/527739

**Authors:** Katie L. Millette, Vincent Fugère, Chloé Debyser, Ariel Greiner, Frédéric J. J. Chain, Andrew Gonzalez

## Abstract

Human impacts on genetic diversity are poorly understood yet critical to biodiversity conservation. We used 175,247 COI sequences collected between 1980 and 2016 to assess the global effects of land use and human density on the intraspecific genetic diversity of 17,082 species of birds, fishes, insects, and mammals. Human impacts on mtDNA diversity were taxon and scale-dependent, and were generally weak or non-significant. Spatial analyses identified weak latitudinal diversity gradients as well as negative effects of human density on insect diversity, and negative effects of intensive land use on fish diversity. The observed effects were predominantly associated with species turnover. Time series analyses found nearly an equal number of positive and negative temporal trends in diversity, resulting in no net monotonic trend in diversity over this time period. Our analyses reveal critical data and theory gaps and call for increased efforts to monitor global genetic diversity.

## Introduction

Intraspecific genetic diversity, a measure of the genetic variation within populations, is an essential measure of biodiversity (Hughes *et al.* 2008). Genetic diversity reflects past and current evolutionary bottlenecks and indicates a population’s potential for adaptation to future stressors (Hewitt 2000; Reed & Frankham 2003; Frankham 2005; Bijlsma & Loeschcke 2012). As such, understanding the drivers of genetic diversity change worldwide is of great interest to ecologists and conservation biologists (Hughes *et al.* 2008; Pereira *et al.* 2013; Mimura *et al.* 2017; Paz-Vinas *et al.* 2018). Human disturbances, acting as an evolutionary force by modifying rates of extinction and colonization (Palumbi 2001; Alberti 2015; Thomas 2015; Schlaepfer *et al.* 2018), may be altering the intraspecific genetic diversity of plants and animals around the world–yet no global assessment of temporal trends in genetic diversity has been conducted to date, nor have human impacts on such trends been quantified.

Theory predicts that human activities can affect intraspecific genetic diversity via demographic and evolutionary mechanisms (Kimura 1968, 1983; King & Jukes 1969). Depending on how human disturbances alter selection, drift, gene flow, and mutation rates, intraspecific genetic diversity may decrease, increase, or remain unchanged over time (DiBattista 2008). For example, habitat fragmentation and harvesting can reduce diversity directly due to high mortality and reduced population sizes, and indirectly due to population isolation and inbreeding (Banks *et al.* 2013). Alternatively, human disturbances can maintain or increase genetic diversity through time, for example by increasing connectivity between populations, magnifying temporal variation in selection, increasing mutation rates (e.g. mutagenic pollutants), or creating environments that favour hybridization and heterozygote advantage (Dubrova *et al.* 1996; Ellegren *et al.* 1997; Bickham *et al.* 2000; Crispo *et al.* 2011).

Trends in intraspecific genetic diversity are expected to be scale-dependent, as are trends in other dimensions of biodiversity like taxonomic, phylogenetic, and functional diversity (McGill *et al.* 2015; Jarzyna & Jetz 2018; Schlaepfer *et al.* 2018; Chase *et al*. 2019). Moreover, human disturbances occurring at different scales may have contrasting effects on genetic diversity. At local scales, high human population density and intensive land use can reduce genetic diversity by lowering habitat quality and population size (Vellend 2004; Coors *et al.* 2009; Helm *et al.* 2009). At regional scales, habitat fragmentation and increased habitat heterogeneity can contribute to genetic differentiation via reduced gene flow (e.g. Ripperger *et al.* 2013) and diversifying selection (e.g. Merck *et al.* 2003), respectively. However, analyses at spatial scales larger than the geographic extent of interbreeding populations will inflate diversity values, particularly when individuals from distant and genetically divergent groups are aggregated. Thus, if the effects of human activity on diversity are to be detected, they should be apparent and strongest at small spatial scales.

Much remains to be learned regarding human impacts on intraspecific genetic diversity. A recent global assessment of mammal and amphibian mtDNA diversity found evidence of reduced diversity in human-impacted regions (Miraldo *et al.* 2016). Given the conservation-relevance of this finding, it is paramount to confirm the robustness of this pattern across spatial scales and taxa while controlling for important confounding variables such as the decay of genetic similarity with distance (Gratton *et al.* 2017) and phylogenetic differences in mtDNA mutation rates (Nabholz *et al.* 2009). Moreover, we are only beginning to understand how human land use is affecting levels of genetic diversity. On a decadal timescale, landscape perturbations would be expected to affect diversity mostly via their effect on population size (Whitlock 1992). However, it remains unclear whether net overall trends in genetic diversity, and human impacts on such trends, exist and can indeed be detected at the global scale.

Here, we evaluate the impacts of human population density and intensive land use on animal intraspecific genetic diversity worldwide, at four spatial scales while accounting for effects of geography, time, and spatial distance between sequences. We use location and time-referenced mitochondrial cytochrome *c* oxidase subunit I (COI) sequences collected between 1980 and 2016 to estimate nucleotide diversity (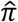), a measure of population genetic diversity, for four animal classes‒birds, inland and coastal bony fishes, insects, and mammals. We test two main predictions: 1) genetic diversity is lower in animal populations exposed to intensive land use and high human population density, and this relationship is strongest at small spatial scales; 2) genetic diversity has declined during the 1980-2016 time-period in highly human-impacted regions.

## Materials and methods

### Sequence data and human impact estimates

Mitochondrial cytochrome *c* oxidase subunit I (mtDNA COI) sequences for birds (Aves), fishes (Actinopterygii), insects (Insecta), and mammals (Mammalia) were downloaded from the National Center for Biotechnology Information (NCBI) ‘GenBank’ (Benson *et al.* 2013) and from the ‘Barcode of Life Data Systems’ (BOLD; Ratnasingham & Hebert 2007) in April 2017. Sequences from GenBank were retrieved with the Entrez Utilities Unix Command Line, and from BOLD using the application platform interface. Only sequences with documented geographic coordinates and sampling dates were downloaded. Sequences with ambiguous taxonomic assignment (e.g. species name containing ‘.spp’, ‘var’) were not considered, nor were the few (< 0.1%) sequences collected before 1980. Species and year-specific sequence alignments were performed using default parameters in MAFFT (Katoh & Standley 2013).

Global human population and land use estimates were obtained from the most recent version of the ‘History Database of the Global Environment’ (HYDE 3.2; Klein Goldewijk *et al.* 2017) for years 1980-2016. HYDE 3.2 provides land use and human population density estimates for all land masses divided into 5 arc-minute grid cells (∼85 km^2^ at the equator). Estimates are available for every decade from 1980 to 2000, and for every year after 2000. HYDE variables included in our analyses were: maximum land area (km^2^/grid cell), human population counts (inhabitants/grid cell), and four categories of intensive land use: cropland, pasture, converted rangeland, and built-up area (km^2^/grid cell). See Klein Goldewijk *et al.* (2017) for a detailed description of these variables. HYDE datasets were converted to raster data structures using the ‘raster’ package (Hijmans 2015) in R version 3.5.0 (R Core Team 2018). Cropland, pasture, converted rangeland, and built-up areas were divided by grid cell land area to estimate the proportion of each cell consisting of the respective category. We then computed two grid cell-specific human impact variables: (1) ‘human population density’ (inhabitants/km^2^), i.e. human population counts divided by grid cell land area, and (2) ‘land use intensity’ (ranging from 0 to 1), i.e. the summed proportions of cropland, pasture, converted rangeland, and built-up area in a grid cell.

All COI sequences were attributed human population density and land use intensity values based on the sequence’s geographic coordinates and the corresponding HYDE grid cell. Sequences sampled between 2000 and 2016 were assigned year-specific HYDE values, while sequences from 1980-89 and 1990-99 were assigned values from the 1980 and 1990 HYDE maps, respectively. Most sequences (92.3%) had year-specific HYDE values (mean +/-SD lag time between year of sequence collection and HYDE 3.2 map = 0.4 +/-1.7 years). Sequences falling on the border of two grid cells, or in cells composed exclusively of water, were excluded. We included data from aquatic animals in grid cells with some land, reasoning that land use can affect inland and coastal waters and the species therein (Stoms *et al.* 2005; Dudgeon *et al.* 2006). All data were processed using the ‘tidyverse’ collection of R packages (Wickham & RStudio 2017). We quantified the proportion of species, genera, families, and orders represented by our final dataset (see ‘Supplementary Methods’) and visualized the distribution of COI sequences according to geography, taxonomy, time, human population density, and land use intensity.

### Calculating genetic diversity and human impacts at the population scale

To split species-specific sequences into distinct geographic populations, we grouped all sequences from a given species across years and mapped their spatial distribution. A simple agglomerative (bottom-up) hierarchical clustering algorithm was used to group sequences based on spatial proximity following the ‘single linkage’ clustering rule (Sneath 1957). Sequences are grouped together as long as the maximum geographical distance among all sequences in the group does not exceed an *a priori* maximum distance threshold (x km). For example, with x = 10 km, two sequences, A and B, would be grouped in the same population if they are 5 km apart, but a third sequence, C, would be placed in its own population if it is >10 km from both A and B. This clustering method was implemented using the single ‘hclust’ function, and a distance matrix of ‘great circle’ geographic distances among sequences with the ‘spDists’ R function (‘sp’ package; Pebesma *et al.* 2018). To assess the scale-dependence of our results, we manipulated ‘x’ in the clustering algorithm to generate new populations with minimum spatial distances of 10, 100, 1,000, and 10,000 km (henceforth: ‘scales’). At large scales, isolated populations may be grouped into a single population, while at small scales, one (true) population may be split into separate populations (see ‘Supplementary Methods’). Without species-specific range and dispersal information, altering the clustering distance criterion was the best approach to assess the scale-dependence of results.

Each sequence was attributed a unique population ID at each spatial scale (e.g. common redpoll, *Acanthis flammea*, from year 2001, in population ‘1’ at scale = 10 km, with mean geographical location 52.4 °N and 4.54°W). Pairwise nucleotide differences were calculated among all sequences in a population with > 50% sequence overlap as in Miraldo *et al.* (2016), using adapted Julia script (https://github.com/mkborregaard/). The mean pairwise nucleotide dissimilarity among sequences was calculated for all populations with > 1 sequence to estimate nucleotide diversity (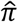; henceforth: ‘diversity’; Nei & Li 1979; Tajima 1993). Diversity was also calculated using stricter inclusion criteria (> 5 sequences; see sensitivity analysis described below). Diversity estimates with values 10 standard deviations greater than the mean of all estimates were discarded. Taxonomic misidentification and/or sequencing error might explain these outliers, which collectively accounted for < 0.1% of the full dataset. For populations with multiple years of sequence data, separate 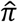 values were computed for each year to assess temporal trends. It should be noted that spatial scale affects the number of sequences and populations in the dataset by affecting both the number of populations into which species-specific sequences are split, and the probability that a sequence is rejected from analysis for being alone in its population.

Following data collection and filtration steps, our dataset consisted of 175,247 COI sequences from 17,082 species of birds (Aves), inland and coastal bony fishes (Actinopterygii), insects (Insecta), and mammals (Mammalia). These sequences were aggregated into 17,124– 27,588 ‘populations’ depending on the spatial scale (Table 1). For each population and at each scale, we calculated the geographical centroid of the population (latitude and longitude), the mean great circle distance among sequences, and the mean land use intensity and human population density of HYDE grid cells from which the sequences originated. Land use heterogeneity within populations could not be incorporated in the analyses because it was strongly collinear with the mean geographic distance among sequences, a well-known predictor of genetic diversity (Gratton *et al.* 2017).

**Table 1.** The number of COI sequences, populations, species, genera, families, and orders included in the dataset of each animal class at each spatial scale.

| scale | class | sequences | populations | species | genera | families | orders |
| --- | --- | --- | --- | --- | --- | --- | --- |
| 10 | Aves | 6155 | 2026 | 1294 (12.50%) | 677 (30.30%) | 120 (52.86%) | 31 (77.50%) |
| 10 | Actinopterygii | 18057 | 4492 | 2297 (7.06%) | 1022 (20.82%) | 240 (49.18%) | 43 (93.48%) |
| 10 | Insecta | 105736 | 18904 | 9714 (1.05%) | 4434 (5.53%) | 430 (38.05%) | 24 (85.71%) |
| 10 | Mammalia | 15632 | 2166 | 531 (9.07%) | 219 (17.03%) | 42 (26.58%) | 10 (34.48%) |
| 100 | Aves | 6834 | 2174 | 1413 (13.64%) | 711 (31.83%) | 120 (52.86%) | 31 (77.50%) |
| 100 | Actinopterygii | 19194 | 3973 | 2366 (7.28%) | 1044 (21.27%) | 245 (50.20%) | 44 (95.65%) |
| 100 | Insecta | 114674 | 17677 | 10740 (1.16%) | 4730 (5.90%) | 442 (39.12%) | 24 (85.71%) |
| 100 | Mammalia | 16248 | 1517 | 547 (9.35%) | 224 (17.42%) | 43 (27.22%) | 11 (37.93%) |
| 1,000 | Aves | 7931 | 1977 | 1561 (15.07%) | 746 (33.39%) | 121 (53.30%) | 31 (77.50%) |
| 1,000 | Actinopterygii | 19919 | 2884 | 2401 (7.38%) | 1054 (21.48%) | 245 (50.20%) | 44 (95.65%) |
| 1,000 | Insecta | 125169 | 14591 | 12006 (1.29%) | 5125 (6.40%) | 452 (40.00%) | 25 (89.29%) |
| 1,000 | Mammalia | 16673 | 743 | 564 (9.64%) | 228 (17.73%) | 43 (27.22%) | 11 (37.93%) |
| 10,000 | Aves | 8597 | 1614 | 1607 (15.52%) | 760 (34.02%) | 123 (54.19%) | 31 (77.50%) |
| 10,000 | Actinopterygii | 20126 | 2427 | 2412 (7.42%) | 1056 (21.52%) | 246 (50.41%) | 44 (95.65%) |
| 10,000 | Insecta | 129705 | 12515 | 12496 (1.35%) | 5261 (6.57%) | 456 (40.35%) | 25 (89.29%) |
| 10,000 | Mammalia | 16819 | 568 | 567 (9.69%) | 228 (17.73%) | 43 (27.22%) | 11 (37.93%) |

Scale indicates maximum geographic distance (km) allowed between any two sequences when
defining populations. Percentages in parentheses for species, genera, families and orders are
estimates of the proportion of taxa present in the database relative to the total number of taxa
globally, as described in Supplementary Methods. Note that the percentage for Actinopterygii
orders is an overestimate caused by large discrepancies in order number in different taxonomic
databases.

### Spatial analyses of animal genetic diversity

Spatial analyses enabled us to test our first prediction that mtDNA diversity is lowest in areas of intensive land use and high human population density, and that this relationship is stronger at small spatial scales. We tested this hypothesis using generalized additive mixed models (GAMMs), a flexible modelling approach, because we did not necessarily expect linear relationships between diversity and human impact variables (e.g. Dornelas *et al.* 2014; Daskalova *et al.* 2019). Models were constructed separately for each animal class and spatial scale, and fitted with the ‘bam’ function in the ‘mgcv’ R package (Wood 2017). All GAMMs used a Tweedie error structure (with a log link function) to account for the zero-inflated, positive, and right-skewed distribution of diversity values. We included sequence number as model weights because nucleotide diversity is likely better estimated with an increasing number of pairwise sequence comparisons. Weights were calculated by dividing the (log-transformed) number of sequences contributing to a diversity estimate by the mean (log-transformed) number of sequences of all diversity estimates from a given taxonomic class and spatial scale. All smooth terms in GAMMs were thin plate regression splines unless noted. The mgcv syntax of all models and their ecological question are described in Supplementary Material (Table S1).

Our spatial GAMMs evaluated the effects of human activity on mtDNA diversity while accounting for spatial and geographical factors known to affect diversity (i.e. spatial distance among sequences, latitude, longitude). Models included the following smooth terms: mean geographic (great circle) distance among sequences (log-transformed), latitude and longitude (a Gaussian process smoothed to account for spatial autocorrelation), absolute latitude, mean human population density (log-transformed), and mean land use intensity (log-transformed). All predictors were rescaled from 0 to 1. Random effects included ‘family’ and ‘order’ to account for phylogenetic influences on 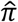, and ‘year of sequence collection’, which could influence diversity estimates if, for example, the error rate of sequencing technology has changed over time. ‘Species’ was not included as a random effect because most species had a single population in the dataset; a single data point (population with the most sequences) was retained instead to avoid non-independence.

Spatial GAMMs were used for two purposes. First, following Gratton *et al.* (2017), fitted values from GAMMs were used to construct global maps of ‘smoothed’ COI nucleotide diversity, after removing the confounding effect of mean spatial distance among sequences in a population (setting this distance to 0 km). Diversity values in 2 x 2° grid cells (a resolution facilitating visualization) were predicted for all combinations of latitudes and longitudes with COI data available. Predicted values were obtained from GAMMs fitted at the 1,000 km scale, which corresponded best to the grain size of the maps. Second, GAMMs were used to visualize and test the significance of relationships between diversity and human variables (human density and land use) after removing spatial effects. We used the ‘summary.gam’ function in mgcv to obtain *F* and *p* values for smooth terms. Predicted relationships between diversity and smooth terms in the model were estimated while setting values for all other predictors to their median value in the taxon- and scale-specific dataset. A condensed summary of model predictions from all 16 GAMMs (4 scales x 4 taxa) is given in the main text. Additional model information can be found in Figs S2‒S5. GAMM validation procedures are described in Supplementary Methods.

We conducted several sensitivity analyses to determine the robustness of our results. First, we performed a rarefaction analysis, refitting all GAMMs after excluding diversity estimates based on < 5 sequences. Second, we refitted models without weights and used the log-transformed number of sequences as a fixed effect/smooth term. Finally, spatial and anthropogenic effects on diversity might differ when considering broad biogeographical regions. Diversity in the Northern and Southern Hemispheres may vary due to differences in geological history (Hewitt 2000, 2004), while dissimilar land uses across biomes (tropical vs. temperate) may bias human effects on diversity. Thus, we fitted additional GAMMs including ‘hemisphere’ or ‘biome’ as factors. None of these analyses resulted in patterns that deviated strongly from the described results (see Supplementary Methods, Results, Figs. S7-12).

Lastly, both species turnover and spatial variation in mtDNA diversity within species may underlie patterns observed in spatial GAMMs. To determine the relative importance of these two processes, we analyzed within-species changes in diversity in the subset of species with data from at least five populations spanning a range of latitudes, human densities, and land use intensities. This analysis focused on the 10 km scale because it provided the most populations and because geographic influences on diversity are minimal at this scale (see Results). For each taxon, we fitted a multi-level GAMM with latitude, human density, and land use intensity as fixed effects, and species as a random effect/smooth, allowing the relationship between diversity and all predictors to vary among species (Table S1). ‘Year of sequence collection’, ‘family’, and ‘order’ were also included as random effects. Predictor variables were centered within species by subtracting, for each population and predictor, their species-specific mean values. This group-mean-centering removes across-species effects when estimating within-species effects of latitude, human density, and land use intensity on diversity.

### Time series analyses

In order to test our second prediction that genetic diversity has declined between 1980 and 2016 in areas of high human impact, we investigated temporal trends in populations that were sampled repeatedly over time (≥ 4 years of data). This analysis was conducted exclusively at the 1,000 km scale because smaller scales produced few time series (< 10 series for birds), whilst at the largest scale (10,000 km), distant individuals could be sampled in different years, conflating spatial and temporal effects. We chose 4 years as the minimum number of years and retained all diversity estimates regardless of their number of sequences because stricter thresholds led to too few time series for reliable analysis. These criteria led to 909 time series from 873 species.

We estimated a Mann-Kendall trend coefficient (τ) for each time series; statistically-significant coefficients indicate decreasing (τ < 0) or increasing (τ > 0) monotonic trends. We also fitted one GAMM per taxon, using a Tweedie error structure and model weights calculated as the mean-standardized number of sequences included in diversity estimates. Fixed effects in these models included latitude, longitude, spatial distance among sequences, year of sequence collection, human population density, land use intensity, and two tensor-product interactions between year and human density, or year and land use intensity (Table S1). These tensor-product interactions allow global temporal trends to vary along anthropogenic gradients, enabling us to test the prediction that diversity is declining in impacted areas. Family and order were included as random effects, as well as a factor-smooth interaction between year and population, to allow population trends to deviate from the global average. We could not fit a random effect for ‘species’ since most species had a single time series, so one series per species (the longest) was retained to avoid non-independence.

## Results

### Data distribution

We visualized the distribution of our sequence data across space, taxonomic groups, time, and anthropogenic gradients. Biases in the spatial distribution of sequences were evident: 70.2% of sequences originated from North America and Europe (Fig. 1a). The dataset was dominated (74%) by insect sequences, while within each class, 1–3 speciose orders contributed a large proportion of sequences (Table 1, Fig. 1b). In all taxonomic classes, COI sequences represented a small proportion of the global number of species, but this proportion increased at higher taxonomic levels (families, orders), suggesting a phylogenetically-broad pattern of sampling (Table 1). Additional information on taxonomic biases can be found in Supplementary Results. The number of sequences collected over time increased steadily in birds, fishes, and insects from 1980 to 2010 (Fig. 1c), while in mammals, the number reached a peak ∼1990 and remained relatively stable until 2005 (Fig. 1c). All groups demonstrate a recent (∼5 year) decline in the number of sequences, perhaps due to the lag between specimen collection and sequence archiving (Fig. 1c). Sequences originated from grid cells generally representative of the global distribution of intensive land use, but from areas with higher human population density than the global average (Fig. 1d, e).

**Figure 1.**
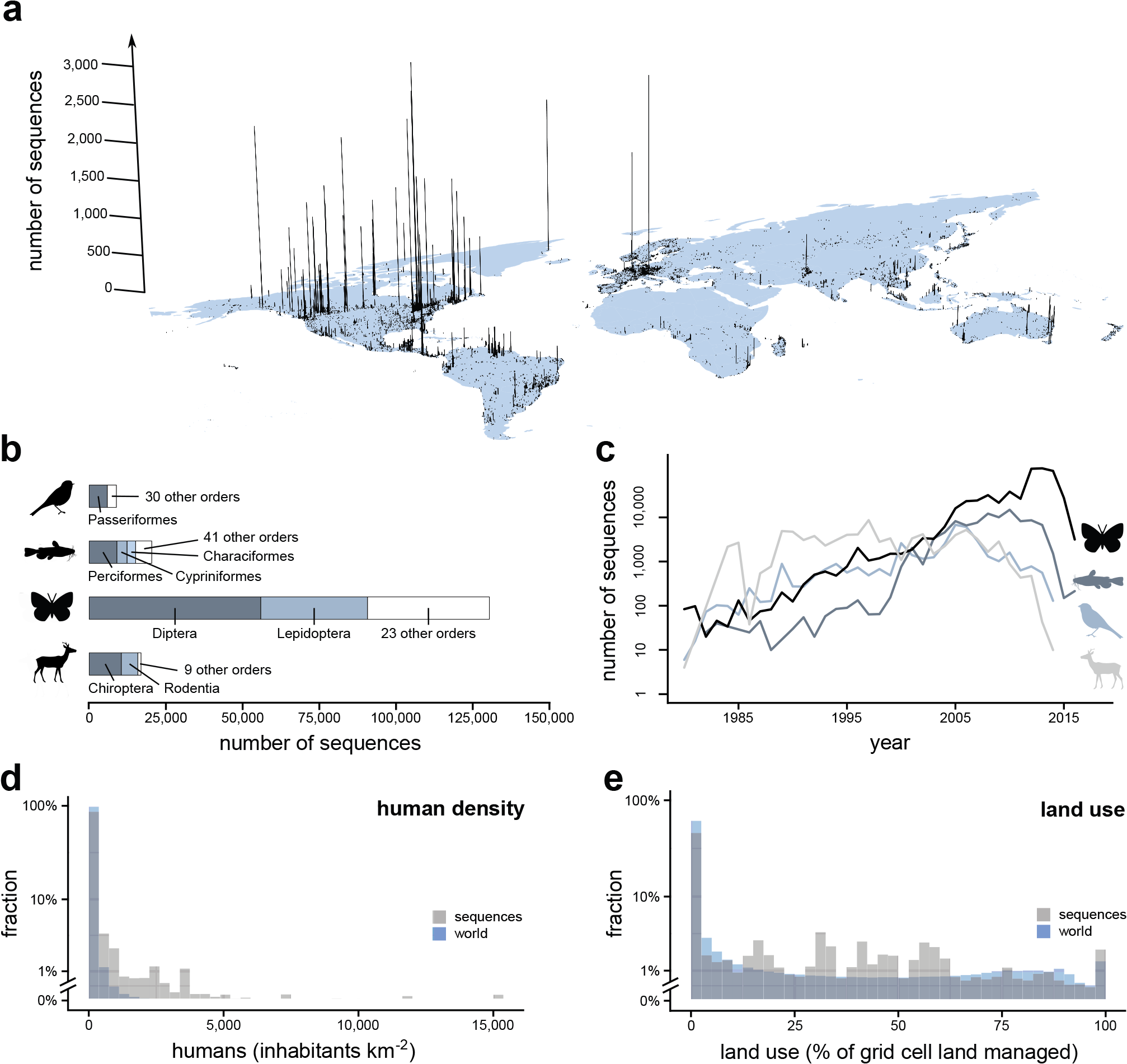
Sequence distribution across geographical space (a), taxonomic classes (b), time (c), and anthropogenic parameters (d, e). COI sequences depicted are those available in NCBI or BOLD with geographic coordinates and year of collection documented in the databases after data filtering steps. (a) Global distribution of sequences. Bar height represents the total number of sequences from the four animal classes per HYDE 3.2 5’ grid cell (min=1 sequence, max=3,258). (b) Distribution of sequences across the four taxonomic classes: birds (N=8,866 sequences), fish (N=20,388), insects (N=130,433), and mammals (N=16,890). Orders contributing ≥10% of a taxon’s sequences are indicated. (c) Number of sequences in the dataset for each year and animal class. (d, e) Distribution of sequences (grey) according to human density (d) and land use intensity (e) relative to the frequency of these parameter values worldwide (blue).

### Spatial analyses of animal genetic diversity

To determine whether mtDNA diversity was lower in areas of high human impact, particularly at small spatial scales, we constructed spatial GAMMs for all combinations of taxon and spatial scale. GAMMs revealed a number of spatial influences on genetic diversity. First, global maps of smoothed COI nucleotide diversity identified a negative latitudinal gradient in birds, fishes, and mammals, but not insects (Fig. 2). Secondly, the mean geographic distance among sequences had a strong positive effect on mtDNA diversity in all taxa (Fig. 3; Figs. S2-S5). However, these spatial effects were scale-dependent: the negative effect of absolute latitude on diversity varied with scale and differed among taxa, and geographic distance was only significant at large spatial scales (Fig. 3). After accounting for spatial effects, we found that human effects on genetic diversity were taxon-dependent and generally weak or non-significant (Fig. 3; Figs. S2-S5). The two clearest effects were: 1) a negative effect of very high human population density (> 5,000 inhabitants/km^2^, or ∼0.75 on the scaled axis) on insect mtDNA diversity at all scales, and 2) the negative impact of heavy land use (grid cells with > 70% intensive land use) on fish mtDNA diversity at all but the largest spatial scale (Fig. 3). A non-monotonic effect of human density on fish genetic diversity was also found at the two largest spatial scales (Fig. 3).

**Figure 2.**
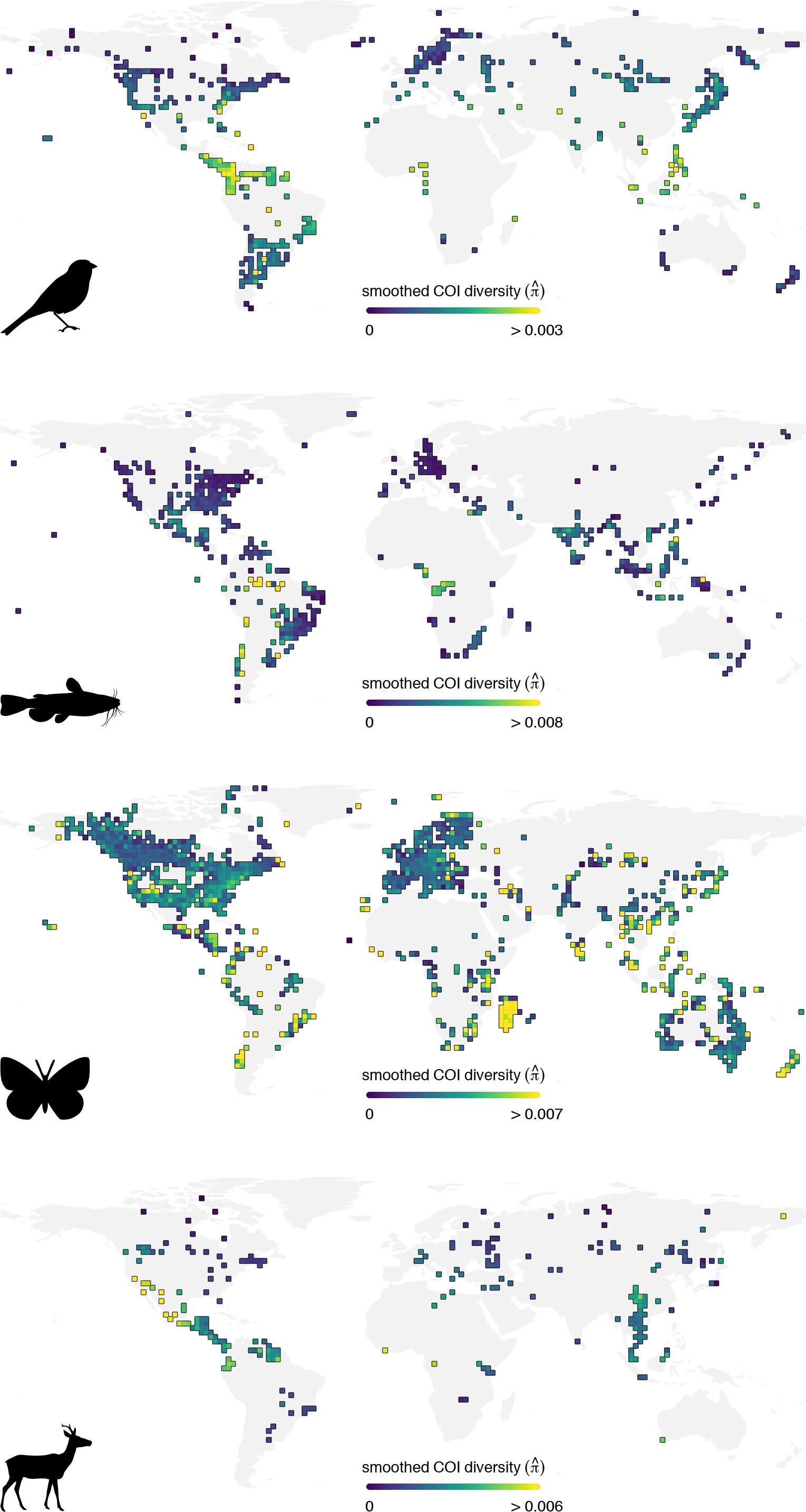
Spatial variation in COI nucleotide diversity of birds, inland and coastal bony fishes, insects, and mammals. Diversity values are GAMM predictions when assuming that all sequences from a given population were collected at the same location, thus removing the confounding effect of population spatial extent on nucleotide diversity. Note that diversity scales differ among animal classes.

**Figure 3.**
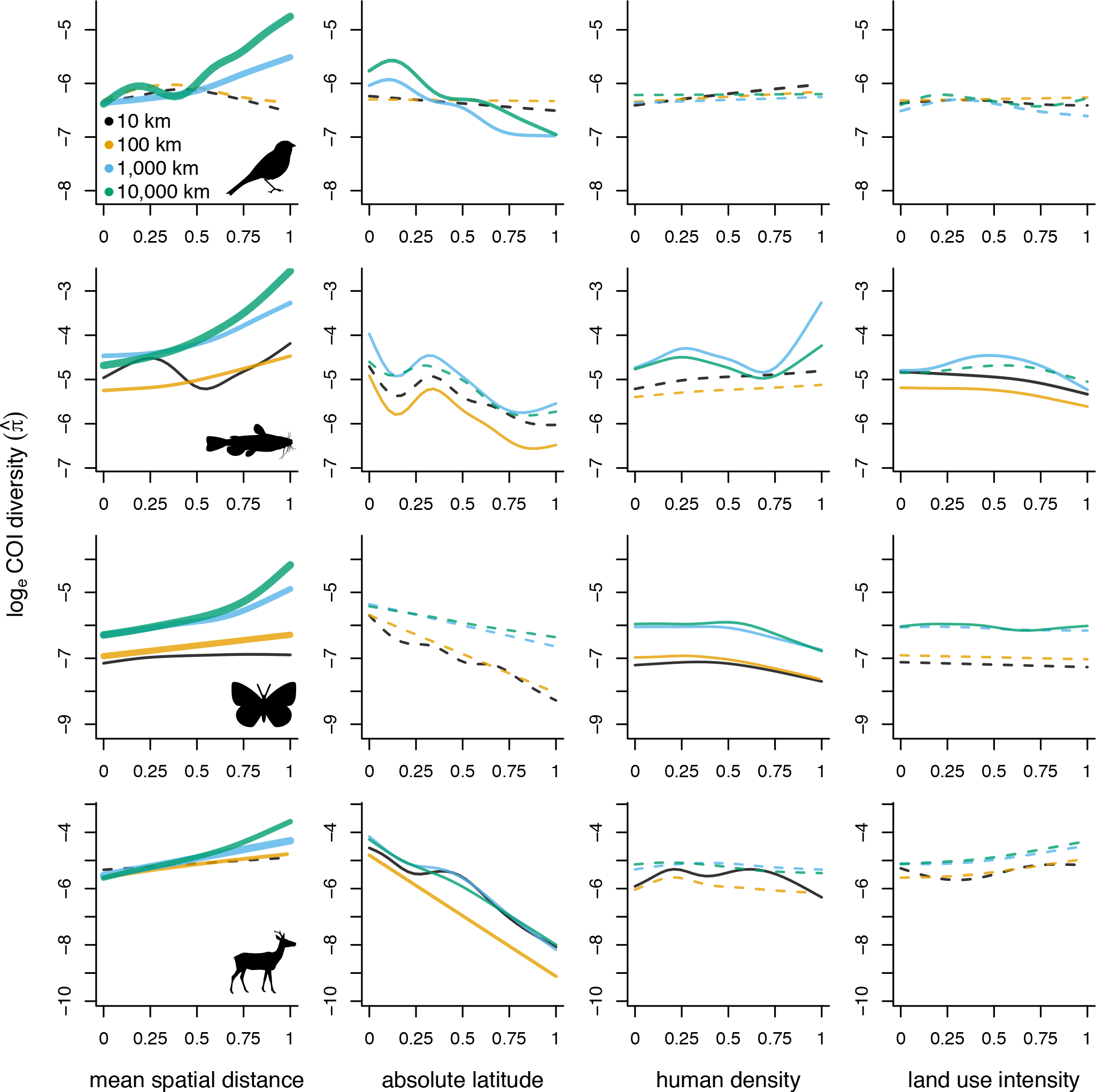
Relationship between COI nucleotide diversity and the mean spatial distance among sequences in a population, the absolute latitude of the population centroid, and the mean human population density, and land use intensity of HYDE 3.2 grid cells from which sequences originate. Panels on the same row belong to the same taxon. Lines represent fitted values from 16 scale and taxon-specific GAMMs (4 scales × 4 classes). Line colour indicates spatial scale; for each taxon, lines with the same colour belong to the same model. Line width indicates effect size (*F* value of predictor) standardized within taxa; thicker lines are stronger effects. Line type indicates statistical significance (solid line, *p* < 0.01; dashed line, *p* > 0.01). Predictor variables were scaled from 0 to 1. Predictions for one variable were made while setting the value of all other variables to their median value in the taxon and scale-specific dataset. Additional model details are provided in Figs. S2-S5.

Rarefaction analysis confirmed the positive effect of geographic distance on genetic diversity at large spatial scales, as well as the negative effects of latitude on mammal mtDNA diversity, of human population density on insect diversity, and of land use on fish diversity (Fig. S7). The latitudinal gradients in birds and fish, and effect of human density on fish genetic diversity however were not robust to rarefaction (Fig. S7). Removing weights from models had little influence on our results (Fig. S8), and incorporating biogeographical factors (biomes, hemispheres) did not reveal clear patterns (Figs. S9-S11), in part due to limited data in the Southern Hemisphere. Analysis of spatial trends in mtDNA diversity within species with multiple population data points suggests that significant effects identified in Fig. 3 are driven primarily by species turnover. After removing species differences in mean latitude, human density, and land use intensity, these predictors had no effect on within-species variation in mtDNA diversity (Fig. 4). Thus, species with relatively higher and lower genetic diversity occur at different locations on latitudinal and anthropogenic gradients, but we find no evidence that these gradients drive spatial variation in genetic diversity within species.

**Figure 4.**
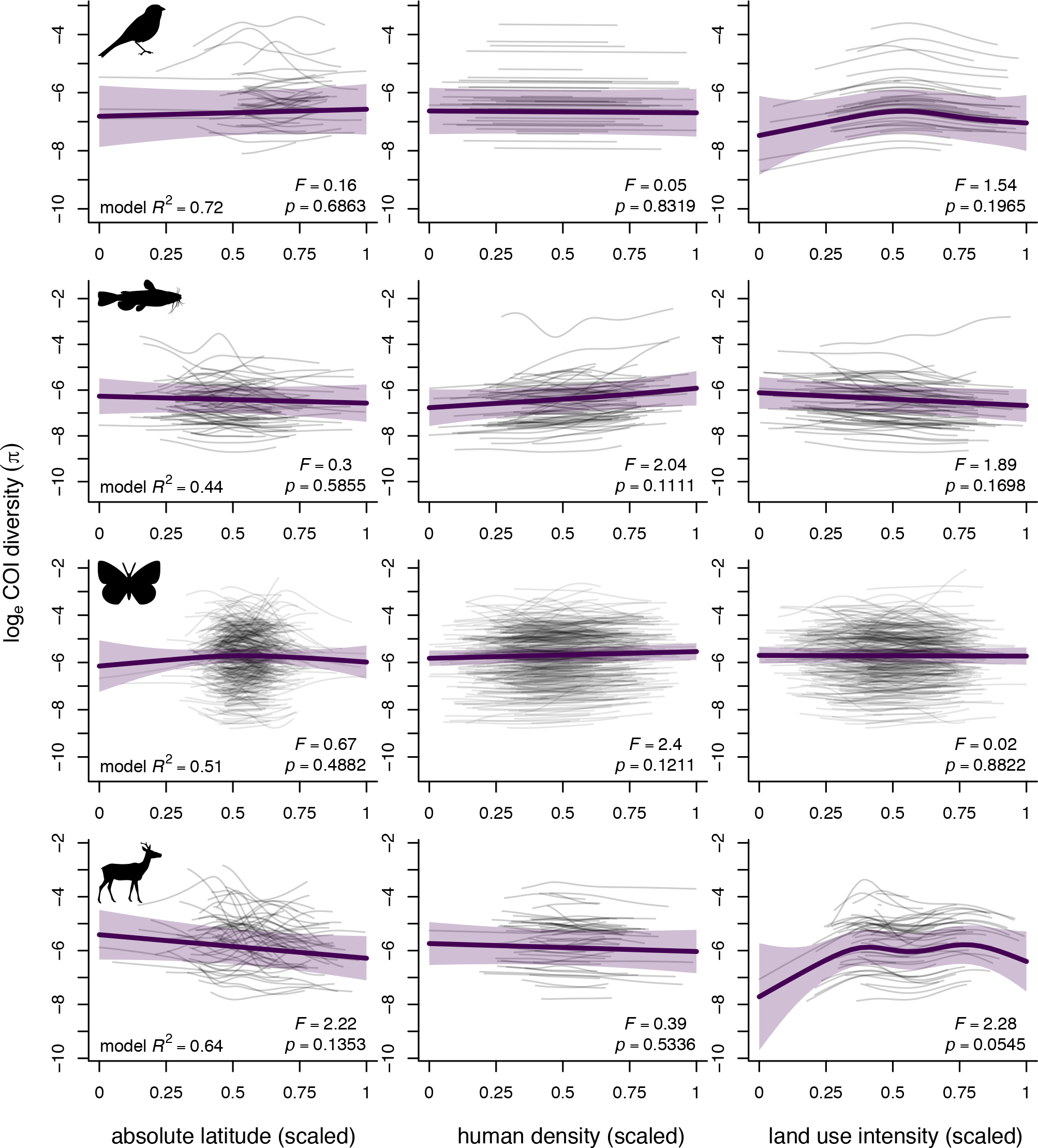
GAMM results showing predicted variation in COI nucleotide diversity within species along gradients of latitude, human population density, and land use intensity. Species included in this analysis had data from ≥ 5 populations varying in their mean values of all three predictor variables at the 10 km spatial scale. Predictor variables were centered using species-specific mean values to ‘align’ all species trends along x axes, thus removing species differences in the mean latitude, human population density, and land use intensity at which they occur. Panels on the same row belong to the same taxon. Thick purple lines indicate overall trends (predicted value ± 95% confidence intervals) and thin grey lines show fitted values for individual species. *F* and *p* values for individual smooths and the overall model adjusted *R*^2^ are shown.

### Time series analyses

To evaluate whether diversity has declined in areas of high human density and land use intensity, we examined temporal changes in mtDNA diversity within populations sampled repeatedly. Most time series showed no significant monotonic trend in diversity, regardless of time series duration (Fig. 5a). In the 2.5% of time series with significant temporal trends, there was nearly an equal number of time series with positive (*N* = 10) and negative (*N* = 12; Fig. 5a) trends. When pooling all populations to estimate a global decadal trend, no significant change in diversity was detected for birds, fishes, and mammals (Fig. 5b). Insects demonstrated a decreasing trend in mtDNA diversity (*p* < 0.01), driven by a modest reduction in diversity from 2010 to 2016 (Fig. 5b; Fig. S12).

**Figure 5.**
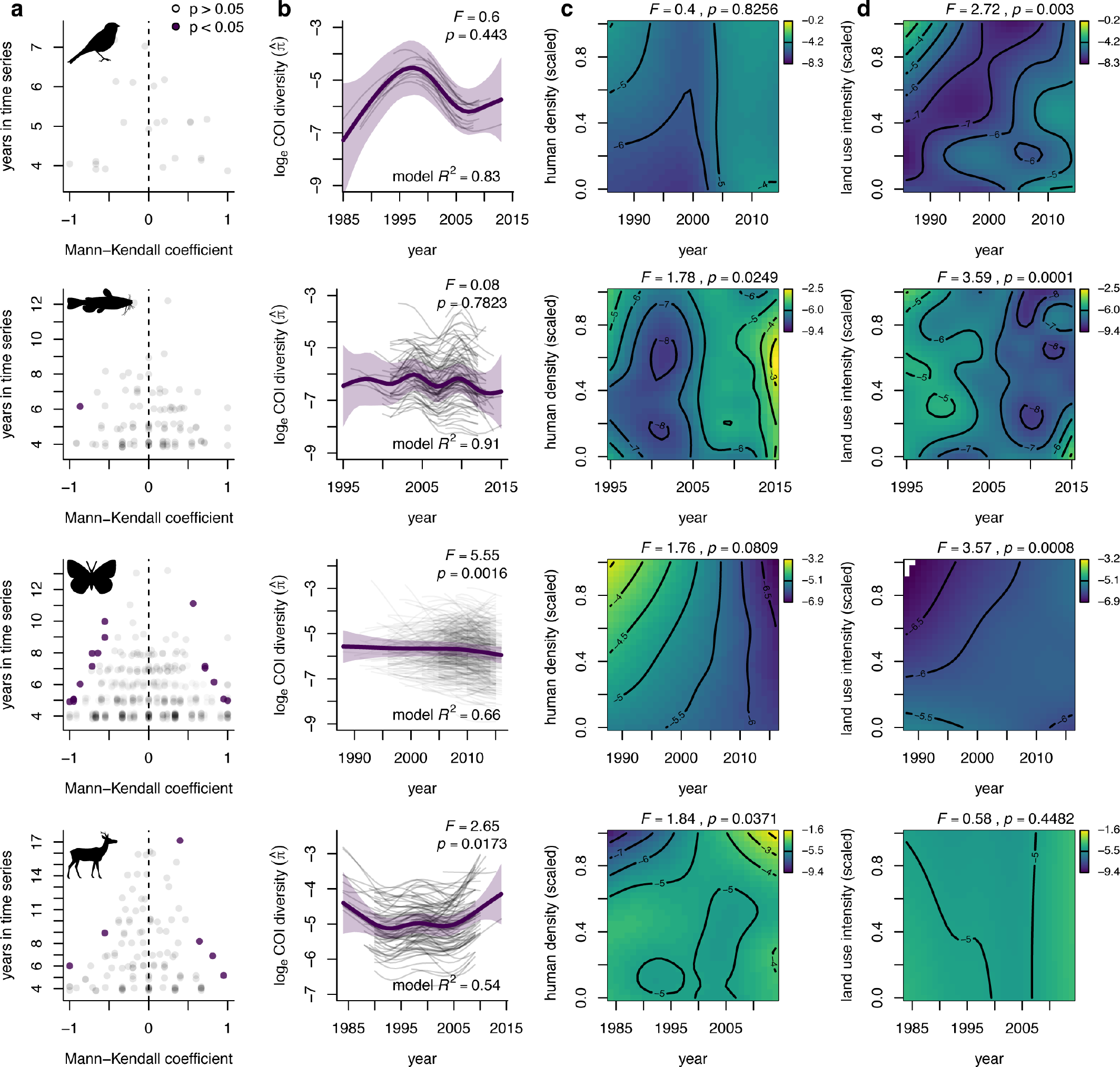
Time series analysis of COI nucleotide diversity in 909 populations of birds, inland and coastal bony fishes, insects, and mammals. Populations were defined at the 1,000 km scale and included a minimum of 4 years of data. Panels on the same row belong to the same taxon. (a) Mann-Kendall trend coefficients (τ) of time series vs. the number of years of data that they contain. Purple symbols are time series with statistically-significant negative (τ < 0) or positive (τ > 0) trends in diversity. A small offset was added on the y axis to aid visualization. (b-d) GAMM results showing predicted values of diversity over time. Values are on the scale of the linear predictor. (b) Effect of year on diversity. Thick purple lines indicate overall trends (predicted value ± 95% confidence intervals) while thin grey lines show fitted values for individual populations. *F* and *p* values for the ‘time’ smooth and the overall model adjusted *R*^2^ are shown. (c, d) Predicted temporal trends in diversity as a function of mean human population density (c) and land use intensity (d) in the HYDE 3.2 grid cells from which population-specific sequences originate. A colour transition from yellow to purple indicates a reduction in nucleotide diversity; for example, in insects (third row, column c), nucleotide diversity in areas of high human population density transitions from high diversity (yellow) to low diversity (purple) over time. The *F* and *p* values of both tensor-product interactions are provided; low *p* values indicate that a given two-dimensional surface is not flat. Note that the scales for diversity and year differ among animal classes.

Several significant interactions between year and human density or land use intensity were found (Figs. 5c, d). This indicates that temporal trends in mtDNA diversity often varied along the two anthropogenic gradients–although effects were generally stronger for land use than for human population density (Figs. 5c, d). Interactions were complex and none were entirely consistent with our prediction of reduced diversity over time in heavily impacted areas. In birds, diversity was highest in areas of intensive land use but declined over time (Fig. 5d). In fishes, an increase in diversity was apparent at intermediate human densities (Fig. 5c), while areas with more intensive land use demonstrated a slightly stronger decline in diversity than areas with less intensive land use (Fig. 5d). Insect mtDNA diversity had a modest temporal increase in areas of intensive land use, while mammal diversity increased over time in areas of high human population densities (Fig. 5c, d).

## Discussion

We found that human density and land use intensity had no clear effect on animal mitochondrial genetic diversity worldwide. In all analyses, the direction and magnitude of human effects were taxon-dependent and either non-significant or weak, even at small spatial scales. We find minimal evidence of a net temporal changes in intraspecific genetic diversity. Time series analyses revealed that the average temporal trend across populations was zero for all taxa but insects, the latter showing a recent, weak, and uncertain decline in mtDNA diversity. Individual populations do demonstrate temporal changes in diversity (2.5% of populations in our dataset), however, with almost an equal proportion of increasing and decreasing trends. Thus, our initial predictions regarding negative effects of humans on animal mtDNA diversity were mostly unsupported, and in contrast with previous work (Miraldo *et al.* 2016).

The geographic distance among sequences was by far the strongest predictor of mtDNA diversity in our global dataset of COI sequences from BOLD and GenBank. This confirms the necessity to account for distance when measuring genetic diversity (Gratton *et al.* 2017), especially when sequences are aggregated across large spatial scales. After accounting for the distance-decay of similarity, latitude still had an influence on the distribution of mtDNA diversity in birds, fishes, and mammals, although this effect was only robust to rarefaction in mammals. This result is consistent with previous studies showing elevated genetic diversity in mammals at low latitudes (Miraldo *et al.* 2016; Gratton *et al.* 2017).

Human impacts on the environment such as urbanization and land use intensification can have both negative and positive effects on intraspecific variation (DiBattista 2008; Hendry *et al.* 2008; Alberti *et al.* 2017; Fugère & Hendry 2018) and other metrics of biodiversity such as species richness (Pautasso 2007; Elahi *et al.* 2015; Daskalova *et al.* 2019). It is therefore not surprising that we find complex and taxon-dependent human effects on mtDNA diversity–that is, it is overly simplistic to expect an overall trend of lower genetic diversity in impacted areas. We did find lower insect genetic diversity in areas with high human densities, which is consistent with recent reports of large declines in insect abundances attributed in part to habitat loss, agriculture, and urbanization (Hallmann *et al.* 2017; Lister & Garcia 2018). Moreover, we also found intensive land use had a negative effect on fish genetic diversity, in both spatial and time series analyses. However, the patterns we report are likely due to species turnover, such that it remains unclear whether species in impacted areas ‘lose’ intraspecific diversity when colonizing these habitats, or whether they have naturally lower diversity than species unable to persist in human-altered environments. These findings stress the need for species-specific monitoring programs across environmental and temporal gradients.

While our results do not support the general conclusion that intraspecific mtDNA diversity is lower in human-dominated areas (Miraldo *et al.* 2016), human effects are currently difficult to evaluate because of several data limitations. Areas with heterogeneous land use may harbour higher genetic diversity than homogeneous landscapes; however, we could not incorporate land use heterogeneity in statistical models due to its strong collinearity with the mean pairwise distance among sequences. Land use estimates from the HYDE datasets are also uncertain due to important model assumptions about mechanisms of land use change (Klein Goldewijk *et al.* 2017). Another limitation is that a large fraction of populations in our dataset originated from a single ‘site’ (geographic coordinates). Without spatial variation among sequences, it is difficult to test the importance of spatial scale and land use on genetic diversity. At large spatial scales (10,000 km), sequences do vary in geographic location, but to the extent that isolation-by-distance becomes the main determinant of genetic diversity. Future assessments of trends in global genetic diversity could incorporate species traits, dispersal potential, phylogeny, and land use history, all of which could influence relationships between diversity and anthropogenic drivers (Wiens 1989; Levin 1992; Chave 2013; Schiebelhut & Dawson 2018; Tucker *et al.* 2018; Binks *et al.* 2019).

We acknowledge that the COI locus does not evolve under neutrality (Pentinsaari *et al.* 2016) and that sequence variation at this mitochondrial locus does not reflect genomic intraspecific diversity, nor may it necessarily reflect anthropogenic pressures (Bazin *et al.* 2006; Nabholz *et al.* 2008). More appropriate genetic metrics are known for measuring neutral intraspecific diversity (e.g. microsatellites, allelic diversity) and are themselves more sensitive to temporal dynamics in ecological processes; however, there is currently no global database for these data (Leigh *et al.* 2019). COI is one of few genes with abundant sequences in common databases (Porter & Hajibabaei 2018) with metadata readily available (e.g. spatial coordinates, date of collection) for many species (likely because COI is used for species identification). Despite the wealth of COI data with collection years in the databases, we noted taxonomic gaps such as the small number of Coleoptera sequences, which as a group account for almost 40% of insect species. As outlined in Supplementary Results, a few species contributed a disproportionately large number of sequences. Moreover, the majority of populations in our dataset were represented by < 10 sequences and mostly from a single year. These data limits hinder the precision of diversity estimates, constrain time series analyses, and impede our ability to assess global patterns of genetic diversity relative to human activity.

In conclusion, anthropogenic activity likely has complex, taxon-specific effects on intraspecific genetic diversity; with the data at hand, one cannot conclude that genetic diversity is generally lower in human-impacted areas. There is a clear need to establish a global and systematic monitoring program to gather more data on intraspecific diversity (Mimura *et al.* 2017). More and longer time series of genetic diversity within populations are urgently needed, as the lack of replication in time is a persistent problem in ecology that is constraining our ability to assess global patterns of biodiversity change (Gonzalez *et al.* 2016; Estes *et al.* 2018; White 2019). Theory should also be developed to determine the conditions under which significant human impacts on genetic diversity are to be expected, as the time scale and demographic consequences of disturbances may not be sufficiently large to affect intraspecific mtDNA diversity (Nabholz *et al.* 2009). Finally, we urge data collectors to upload metadata such as collection year and spatial coordinates when depositing sequences in databases–a remarkably large number of sequences in GenBank do not have a collection year (e.g. 95% of amphibian sequences; Marques *et al.* 2013; Pope *et al.* 2015). Global monitoring of genetic diversity would improve our ability to detect change and attribute the causes of worldwide patterns of spatial and temporal variation in genetic diversity.

## Acknowledgements

This work resulted from insightful discussions among past and present Gonzalez lab members. We are grateful to Andreia Miraldo and Michael Borregaard for assistance with Julia scripts. We thank Navin Ramankutty, Erle Ellis, and Kees Klein Goldewijk for help accessing and interpreting the HYDE 3.2 dataset. Simon Joly, Arne Mooers, and Daniel Schoen contributed insightful comments on earlier versions of the manuscript. Additional comments from three anonymous reviewers greatly improved this manuscript. The authors acknowledge support from the Natural Sciences and Engineering Research Council of Canada (NSERC) and the Quebec Centre for Biodiversity Science (QCBS). AG is supported by the Liber Ero Chair in Biodiversity Conservation and a Killam Fellowship.

