## Supporting information for "No consistent effects of humans on animal genetic diversity worldwide"

**This file includes:**

Supplementary Methods
Supplementary Results
Supplementary References
Table S1
Figures S1 to S12

### Supplementary Methods

#### *Spatial and taxonomic coverage*

To visualize our dataset, we examined the distribution of COI sequences with respect to geography, taxonomy, time, and human impact variables (Figure 1 of the main text). We first generated a global map showing the number of sequences falling into each HYDE (5') grid cell, using R packages 'latticeExtra' (Sarkar & Andrews 2016) and 'rworldmap' (South 2011). The number of sequences per year were tallied for each animal class (birds, fishes, insects, and mammals). We also determined whether sequences were under levels of human influence representative of global averages by comparing the distribution of human density and land use intensity values associated with 1) all HYDE 3.2 grid cells, worldwide and pooled across all years, and 2) the time and place of sequence collection.

For each animal class, we also quantified the proportion of global taxa represented in our sequence datasets. We first retrieved genus, family, and order-level classification for all species using taxonomic information from the NCBI and the 'Integrated Taxonomic Information System' (ITIS) databases accessed through the R package 'taxize' (Chamberlain & Szöcs 2013). Supplemental information regarding Actinopterygii order classification was obtained from 'Fishbase' (Froese & Pauly 2018; *www.fishbase.org*) using 'rfishbase' (Boettiger *et al.* 2012). Taxonomic information for species missing full classifications in NCBI, ITIS, and Fishbase (such as obsolete, or synonymous names) was retrieved from BOLD itself. This order was followed to ensure classifications were as current as possible. The total global number of genera, families, and orders in each class was then obtained from the 'Catalogue of Life' 2018 database (Roskov *et al.* 2018; *www.catalogueoflife.org/annual-checklist/2018/*) and compared to the number of taxa in our dataset. This provided a rough estimate of the proportion of taxa included in our dataset; exact proportions cannot be calculated due to unresolved taxonomy and discrepancies among references (Guralnick *et al.* 2007). The proportion of taxonomic representation obtained for Actinopterygii orders (95%) is likely an over-estimate due to large discrepancies in the number of orders present in different databases; for example, some orders represented in BOLD/NCBI (and thus in our dataset) are not present in either Fishbase, nor the Catalogue of Life, and vice versa. The proportion of species representation for birds, insects, and mammals should be more accurate as mismatches across databases were limited.

*Calculating population genetic diversity at various spatial scales*

To assess the scale-dependence of our results, we manipulated the minimum spatial distances in the agglomerative clustering algorithm, using four different minimum spatial distances for the creation of a new population: 10, 100, 1,000, and 10,000 km ('scales'). Depending on the species, at the largest spatial scale (10,000 km), sequences spanning an entire continent may be grouped into one population, likely aggregating sequences from isolated populations. Likewise, at the smallest spatial scale (10 km), sequences from one (true) population may be split into separate populations if no individuals were sampled in the middle of the population's range. Using species-specific dispersal range information to group sequences might be more appropriate; however such information was not available for a majority of species. Moreover, spatial biases in the distribution of COI sequences suggest that population ranges were rarely sampled in their entirety: the mean great circle distance among sequences was 0 for ~50 % of species in our dataset, and usually < 100 km for other populations (Fig. S1). Therefore, with the data at hand, altering the choice of clustering distance criterion was the best compromise to assess the scale-dependence of results.

*Generalized additive mixed model (GAMM) validation and sensitivity analyses*

All models were validated with plots of residuals against fitted values and predictor variables, with autocorrelation functions (for temporal autocorrelation), and with variograms and maps of residuals (for spatial autocorrelation). We ensured that the choice of basis dimension (*k*) was sufficiently high using the ‘gam.check’ function in mgcv. We also conducted a sensitivity analysis to determine whether the low number of sequences contributing to most diversity estimates influenced our results. Indeed, the median number of sequences compared across all diversity estimates was 3 (range = 2–1,434 sequences; Fig. S6). Using sequence number as weights in GAMMs partly accounts for this problem; however, we also re-ran all analyses using a stricter data inclusion criterion, whereby all diversity estimates with fewer than 5 sequences were discarded. Given the dataset, this is the strictest criterion we could apply while keeping the same GAMM structure and fitting models for all combinations of animal classes and spatial scales. Results of GAMMs including only diversity estimates with > 5 sequences are presented in Figure S7. Then, we also determined whether using model weights influenced our results. We refitted all 16 spatial GAMMs without weights but including an additional smooth term: the (log-transformed) number of sequences used to compute diversity estimates. The results of these (unweighted) models are presented in Figure S8. Time series GAMMs (Fig. 5 of the main text) were also validated using ‘gam.check’ and by inspecting model residuals. However, we could not conduct a sensitivity analysis with a stricter data inclusion criterion (e.g. > 5 sequences) because the number of time series was already low with the most permissive criterion (2 sequences or more), especially for birds. We nonetheless refitted all models without weights to verify that weighting did not affect our results. Time series GAMMs without weights but with an extra smooth term (the log-transformed number of sequences contributing to diversity estimates) are shown in Figure S12.

*Biogeographical patterns in spatial GAMMs*

We considered two biogeographical factors which might modulate the trends that we report in spatial GAMMs. More specifically, we hypothesized that: 1) latitudinal diversity gradients might differ between the Southern and Northern Hemispheres due to their different geological histories, and 2) human impacts on diversity might differ between the temperate and tropical biomes because of the different types and proportions of land use in these two areas. To test these hypotheses, we refitted spatial GAMMs while adding an effect of ‘hemisphere’ or ‘biome’. One model included hemisphere (north vs. south) as a parametric effect and a factor-smooth interaction between hemisphere and latitude (using the ‘by’ argument when specifying smooth terms in mgcv); the other model included ‘biome’ (tropical vs. temperate) as a parametric effect and two factor-smooth interactions, between biome and human population density and between biome and land use intensity. Models including both factors (hemisphere and biome) and/or additional interactions did not converge, in part because data for the Southern Hemisphere is limited. To ensure that latitude range would be similar for the Southern and Northern hemispheres, populations from latitudes > 60˚N were excluded, as no populations were available from Southern polar regions. Thus, when testing effects of ‘biome’, we only included populations found between -30 and 30˚ latitude (‘tropical’), and between (-60, -30˚) and (30, 60˚; ‘temperate’). To test whether smooths from the two hemispheres or biomes were significantly different from one another, ‘hemisphere’ and ‘biome’ were coded as ordered factors, leading mgcv to estimate a smooth for the ‘reference’ level (e.g., ‘Southern’ or ‘tropical’) and a *difference smooth* representing the deviation of the other factor level (Northern’ or ‘temperate’) from the reference smooth. The $p$ value of the difference smooth provided by the ‘summary.gam’ function then determines whether smooth terms (absolute latitude, human population density, or land use intensity, depending on the model) are significantly-different between the two factor levels.

### Supplementary Results

*Taxonomic biases*

The bird dataset consists of 66.7% Passeriformes sequences, with a single species (Tenerife blue chaffinch, *Fringilla teydea*), endemic to the Canary Islands, contributing the greatest number of sequences (~1.9% in 6 years) of all bird species. No species dominated the fish dataset, although three speciose orders (Perciformes, Cypriniformes, and Characiformes) contributed about 74% of sequences. Diptera (43%) and Lepidoptera (27%) sequences dominated the insect dataset, with an agricultural pest, commonly known as the seedcorn maggot (*Delia platura*), overwhelmingly abundant at approximately 2% of sequences (collected over 7 years throughout the Northern Hemisphere). Coleoptera sequences were relatively rare (7.6%) given the proportion of insect species that belong to this order. Mammals were overrepresented by bat species (62% Chiroptera), with approximately 12.4% belonging to what appears to be a yearly, long-term (1989-2010) survey of Seba’s short-tailed bat (*Carollia perspicillata*), great fruit-eating bat (*Artibeus lituratus*), and dark fruit-eating bat (*A. obscurus*) in Central and South America.

*Spatial GAMMs*

Spatial GAMMs had low adjusted $R^2$ (< 0.2; Figs. S2-S5), indicating that despite the presence of some significant effects, the predictor variables included in this analysis (latitude, longitude, spatial distance among sequences, human population density, land use intensity, and year of sequence collection) were not, collectively, strong drivers of COI nucleotide diversity. This could in part be due to the low precision of diversity estimates, many of which were based on a handful of sequence comparisons (Fig. S6). When fitting spatial GAMMs with more stringent data inclusion criteria (diversity estimates based on ≥ 5 sequences, giving 10 potential pairwise comparisons or more), we confirmed the strong positive effect of geographic distance among sequences on genetic diversity, apparent in all taxa at the two largest spatial scales (Fig. S7). However, effects of absolute latitude on diversity disappeared (or changed in form) in birds and fishes, and only remained in mammals (Fig. S7), suggesting that a clear latitudinal gradient is only present in this taxon. Negative impacts of human population density on insect genetic diversity and of land use on fish genetic diversity remained apparent in this analysis (Fig. S7).

*Biogeographical patterns in spatial GAMMs: effects of hemisphere and biome*

Latitudinal diversity gradients did not differ markedly between the Southern and Northern Hemispheres (Fig. S9); however, data for the Southern Hemisphere were few, especially at latitudes lower than 45˚ South. Thus, confidence intervals in these models were rather broad. In birds, fishes, and insects, the few significant differences among hemispheres were due to a more 'wiggly' trend for the Southern Hemisphere, and to the large model uncertainty at high latitudes sometimes causing an upward spike in diversity at latitudes exceeding 45˚S. The dominant trend was for a negative effect of absolute latitude in both hemispheres. For mammals, too few data were available from the Southern Hemisphere for model convergence and GAMMs provided nonsensical results (Fig. S9).

When comparing the effect of human population density and land use intensity on diversity in tropical vs. temperate populations, we found few significant differences among biomes (Figs. S10-S11). In birds, no difference was found across biomes for either human population density (Fig. S10) nor land use intensity (Fig. S11). In fishes, very high human population density (> 0.75 on the scaled axis) had opposing effects on diversity in the temperate vs. tropical biome, with a negative effect on diversity in the temperate biome but a positive effect in the tropics (Fig. S10). In insects, high human population density had a clear negative effect on diversity (as reported in the main text), but this effect was more pronounced in the temperate than tropical biome (Fig. S10). In mammals, land use had a negative effect on diversity in the temperate biome but a complex, non-monotonic effect on diversity in the tropical biome (Fig. S11). Note, however, that all of these effects were scale-dependent and weak, with 95% confidence bands overlapping for the two biomes across the whole range of human population density and land use intensity values, for most combinations of scale and taxon.

*Table S1.* Ecological question (Q) and mgcv model formula (F; where applicable) for all statistical analyses described in the main text and in Supplementary Material.

**Abbreviations**

*Variables.* div: genetic diversity. lat: latitude. long: longitude. D: mean geographic distance among sequences forming a population (log-transformed). lat.abs: absolute value of latitude. hd: human population density (log-transformed). p.lu: land use intensity (log-transformed).

*Smooth term basis/types.* gp: Gaussian process. tp: thin plate regression splines. re: random intercept. fs: random smooths. ti: tensor interactions.

**Spatial Analyses**

*Spatial GAMMs with the full dataset (Fig. 3). All spatial scales.*

**Q**: When controlling for spatial distance among sequences in a population and latitude, does genetic diversity vary with human population density and land use intensity?

**F**: div ~ s(lat,long, k=50, bs='gp') + s(D, k=8, bs='tp') + s(lat.abs, k=8, bs='tp') + s(hd, k=8, bs ='tp') + s(p.lu, k=8, bs='tp') + s(year, bs='re', k=5, m=1) + s(order, bs='re', k=5, m=1) + s(family, bs='re', k=5, m=1)

*Spatial GAMMs with hemisphere as an ordered factor (Fig. S9). All spatial scales.*

**Q**: When controlling for spatial distance among sequences in a population, human population density, and land use intensity, is the relationship between latitude and diversity different between the southern vs. northern hemisphere?

**F**: div ~ hemisphere + s(lat,long, bs='gp') + s(D, k = 8, bs = 'tp') + s(lat.abs, k=8, bs='tp') + s(lat.abs, k=8, by=hemisphere) + s(hd, k=8, bs ='tp') + s(p.lu, k=8, bs='tp') + s(year, bs='re', k=5, m=1) + s(order, bs='re', k=5, m=1) + s(family, bs='re', k=5, m=1)

*Spatial GAMMs with biome as an ordered factor (Figs. S10-S11). All spatial scales.*

**Q**: When controlling for spatial distance among sequences in a population and latitude, does genetic diversity vary with human population density and land use intensity, and are the relationships between diversity and human impact variables different between the temperate vs. tropical biome?

**F**: div ~ biome + s(lat,long, bs='gp') + s(D, k=8, bs='tp') + s(lat.abs, k=8, bs='tp') + s(hd, k=8, bs='tp') + s(hd, k=8, by=biome) + s(p.lu, k=8, bs='tp') + s(p.lu, k=8, by=biome) + s(year, bs='re', k=5, m=1) + s(order, bs='re', k=5, m=1) + s(family, bs='re', k=5, m=1)

*Spatial GAMMs with the subset of species that have at least 5 populations that vary in absolute latitude, human population density, and land use intensity (Fig. 4). Scale = 10 km.*

**Q**: When removing species differences in the mean latitude, human population density, and land use intensity at which they occur (with group-mean-centering), does spatial variation in genetic diversity WITHIN species correlates with variation in latitude or human impacts?

**F**: div ~ s(lat.abs, k=8, bs='tp') + s(hd, k=8, bs='tp') + s(p.lu, k=8, bs='tp') + (lat.abs, species, bs='fs', k=6, m=1) + s(hd, species, bs='fs', k=6, m=1) + s(p.lu, species, bs='fs', k=6, m=1 + s(year, bs='re', k=5, m=1) + s(order, bs='re', k=5, m=1) + s(family, bs='re', k=5, m=1)

**Time series analysis**

*Mann-Kendall test with populations sampled at least 4 times (Fig. 5a). Scale = 1000 km.*

**Q**: How many individual populations show evidence of a temporal trend in diversity (increasing or decreasing)?

*GAMMs with populations sampled at least 4 times (Fig. 5b-d). Scale = 1000 km.*

**Q**: Has global diversity been increasing or decreasing over the past decades? Are such global trends different in areas of high vs. low human impacts?

**F**: div ~ s(lat,long, k=50, bs='gp') + s(D, k=8, bs='tp') + s(year, k=8, bs='tp') + s(hd, k=8, bs ='tp') + s(p.lu, k=8, bs='tp') + ti(year, hd, k=8) + ti(year, p.lu, k=8) + s(year, pop, bs='fs', k=5, m=1) + s(order, bs='re', k=5, m=1) + s(family, bs='re', k=5, m=1)

*Figure S1*. Mean Great Circle distance among conspecific sequences (pooling sequences from all years) for all species included in the dataset.

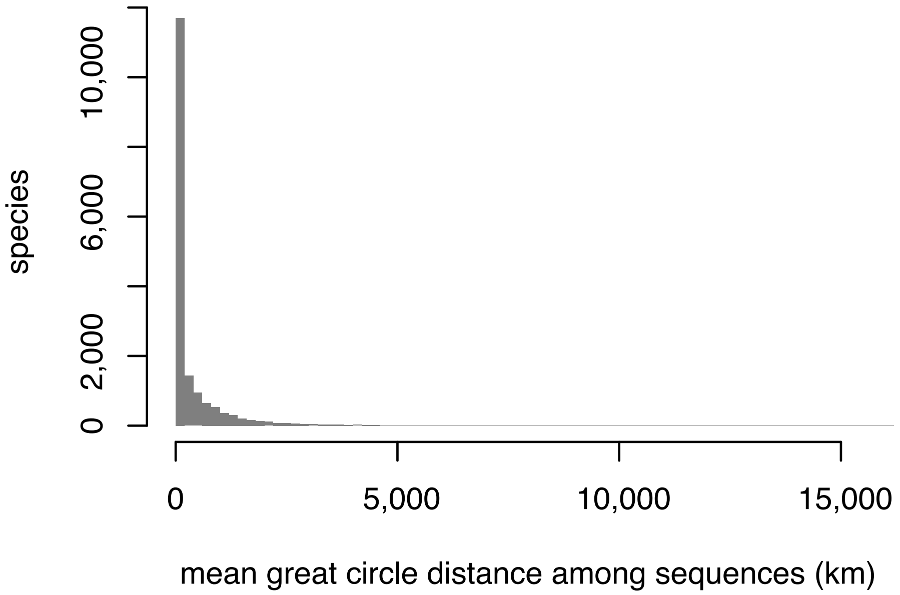

*Figure S2.* Results of the four scale-specific spatial GAMMs constructed for birds, using all diversity estimates (including those with only 2 sequences). Each row of four panels illustrate the results of one GAMM. Line colours match those of Fig. 3 in the main text. For each of the 4 predictor variables of interest, fitted values with 95 % confidence intervals are shown along with the *F* and *p* value of the smooth term given by summary.gam() in mgcv. The overall model $R^2$ is also shown. Predictor variables were scaled from 0 to 1. Predictions for one variable were made while setting the value of all other variables to their median value in the taxon and scale-specific dataset used to fit the GAMM. For visualization purposes, fitted values are shown on the scale of the linear predictor.

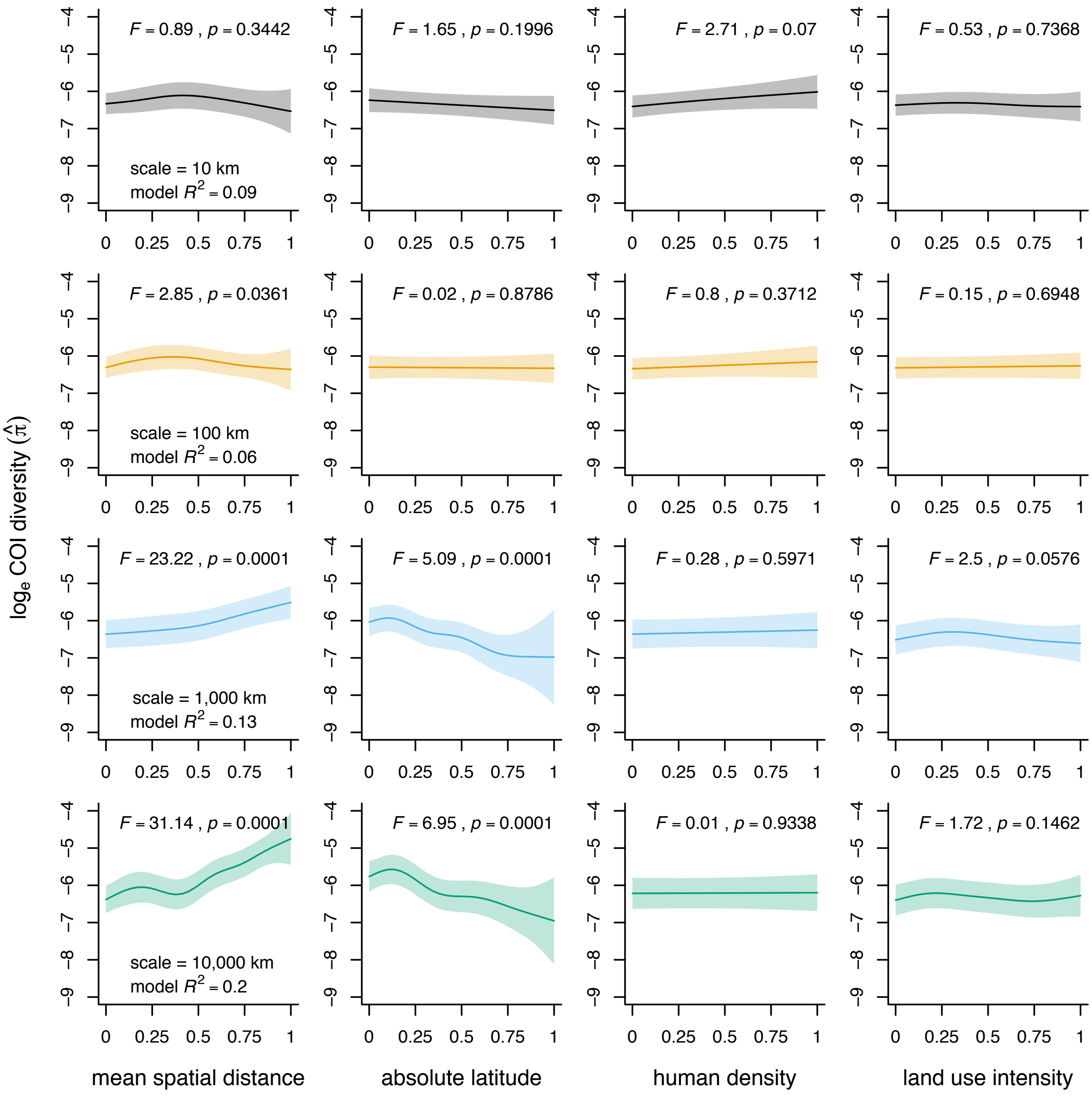

*Figure S3.* Results of the four scale-specific spatial GAMMs constructed for fishes, using all diversity estimates (including those with only 2 sequences). Each row of four panels illustrate the results of one GAMM. Line colours match those of Fig. 3 in the main text. For each of the 4 predictor variables of interest, fitted values with 95 % confidence intervals are shown along with the *F* and *p* value of the smooth term given by summary.gam() in mgcv. The overall model $R^2$ is also shown. Predictor variables were scaled from 0 to 1. Predictions for one variable were made while setting the value of all other variables to their median value in the taxon and scale-specific dataset used to fit the GAMM. For visualization purposes, fitted values are shown on the scale of the linear predictor.

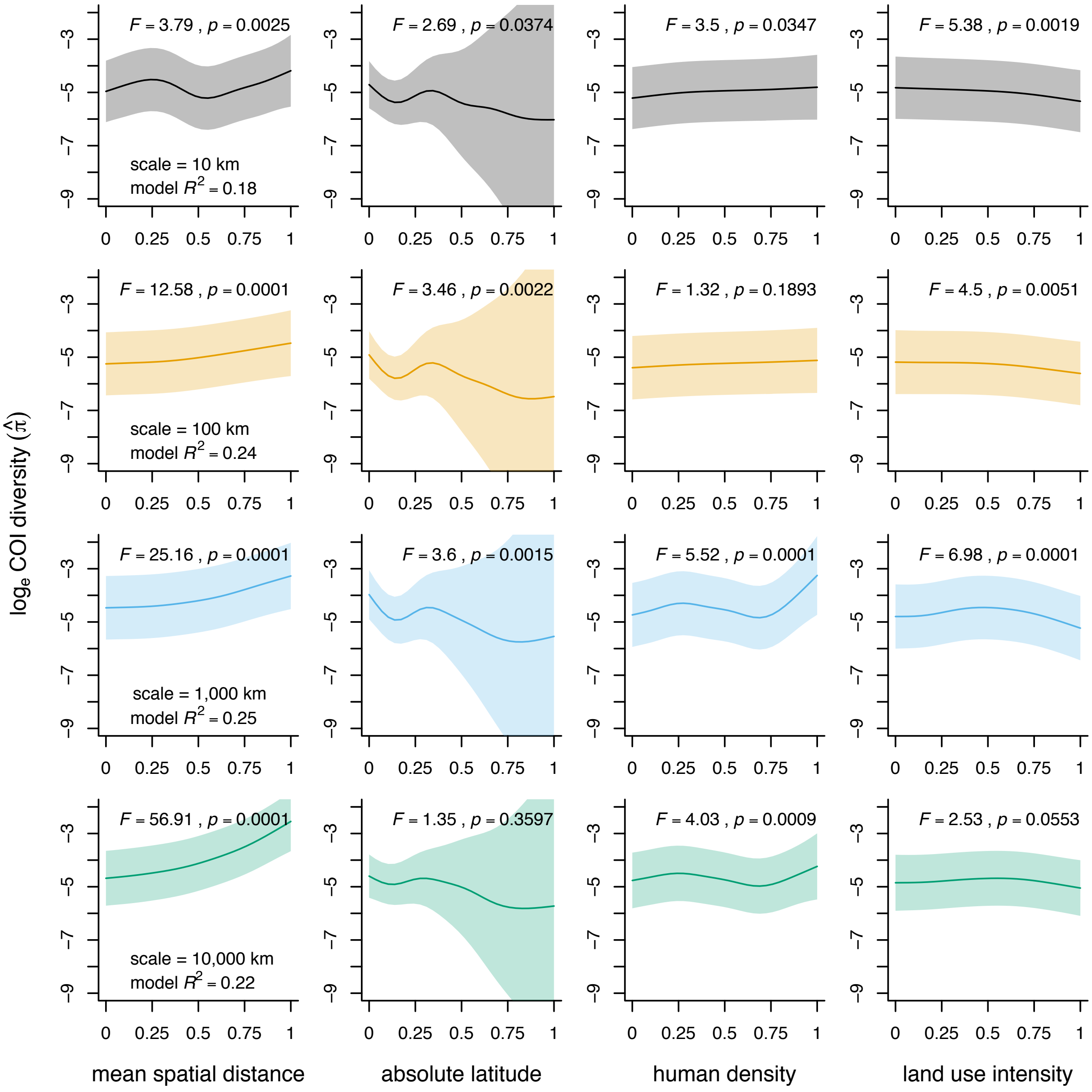

*Figure S4.* Results of the four scale-specific spatial GAMMs constructed for insects, using all diversity estimates (including those with only 2 sequences). Each row of four panels illustrate the results of one GAMM. Line colours match those of Fig. 3 in the main text. For each of the 4 predictor variables of interest, fitted values with 95 % confidence intervals are shown along with the *F* and *p* value of the smooth term given by summary.gam() in mgcv. The overall model $R^2$ is also shown. Predictor variables were scaled from 0 to 1. Predictions for one variable were made while setting the value of all other variables to their median value in the taxon and scale-specific dataset used to fit the GAMM. For visualization purposes, fitted values are shown on the scale of the linear predictor.

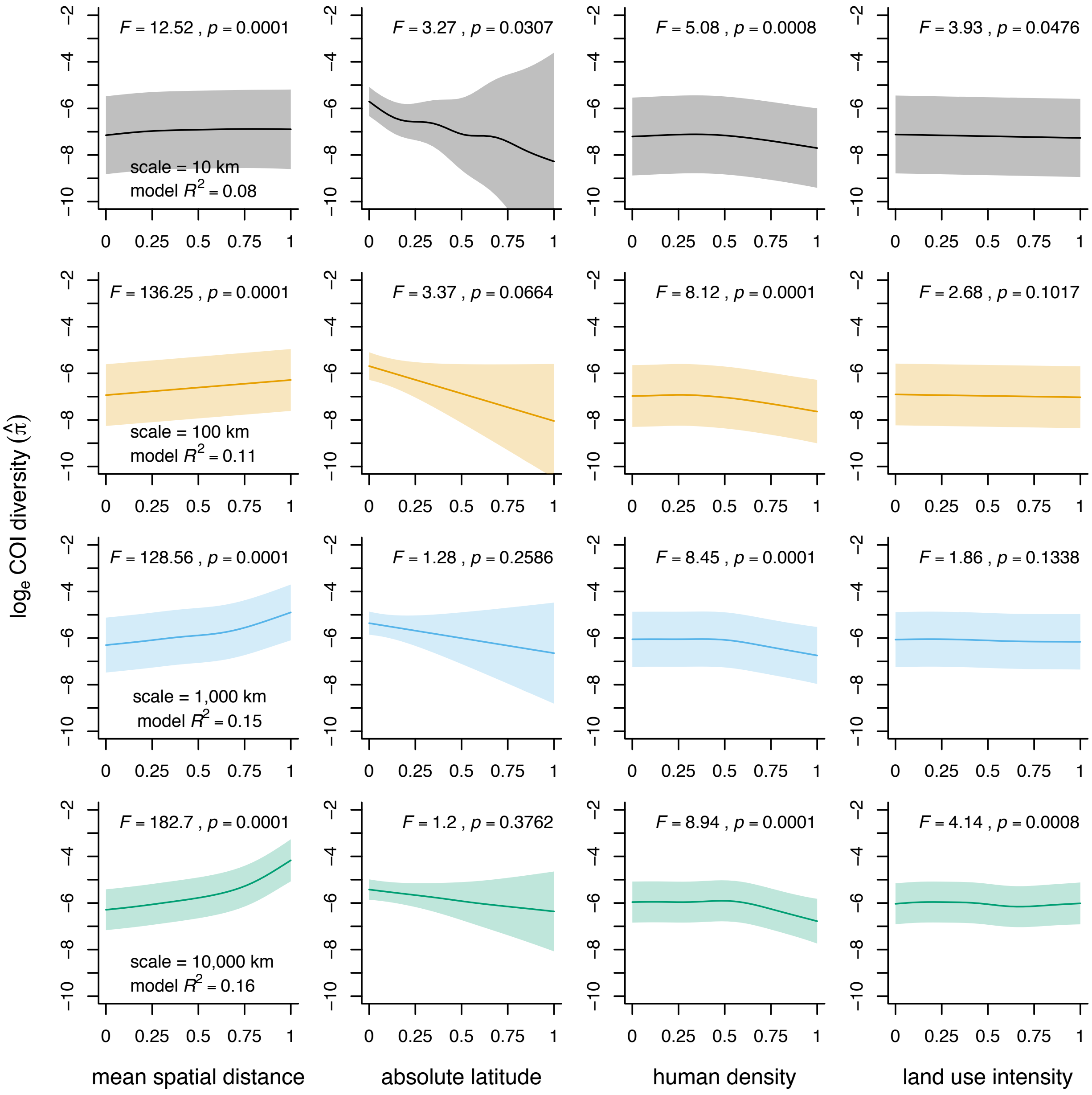

*Figure S5.* Results of the four scale-specific spatial GAMMs constructed for mammals, using all diversity estimates (including those with only 2 sequences). Each row of four panels illustrate the results of one GAMM. Line colours match those of Fig. 3 in the main text. For each of the 4 predictor variables of interest, fitted values with 95 % confidence intervals are shown along with the *F* and *p* value of the smooth term given by summary.gam() in mgcv. The overall model $R^2$ is also shown. Predictor variables were scaled from 0 to 1. Predictions for one variable were made while setting the value of all other variables to their median value in the taxon and scale-specific dataset used to fit the GAMM. For visualization purposes, fitted values are shown on the scale of the linear predictor.

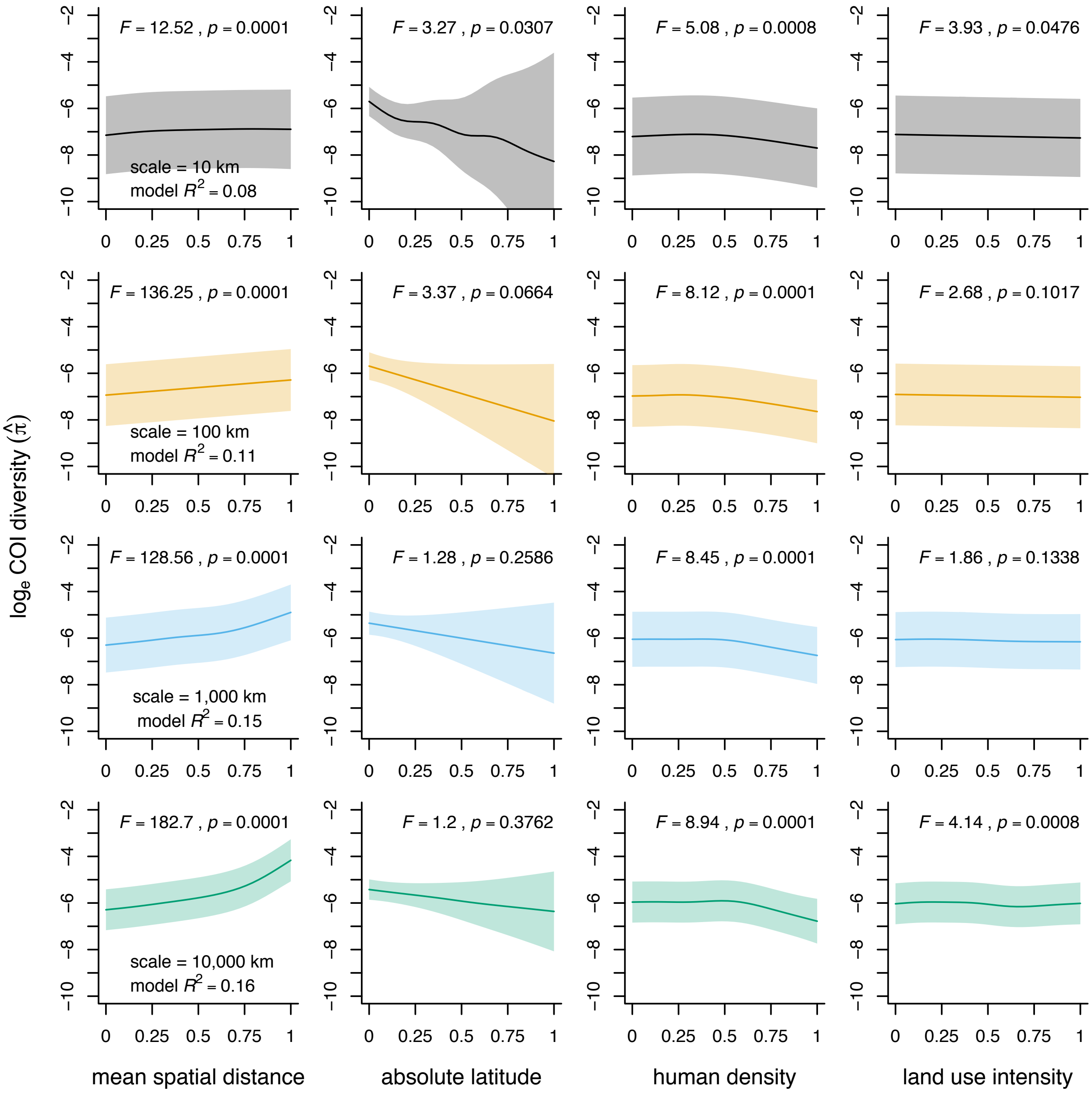

*Figure S6.* Number of sequences compared to calculate nucleotide diversity, for all diversity estimates in the dataset. The x-axis is logged for clarity.

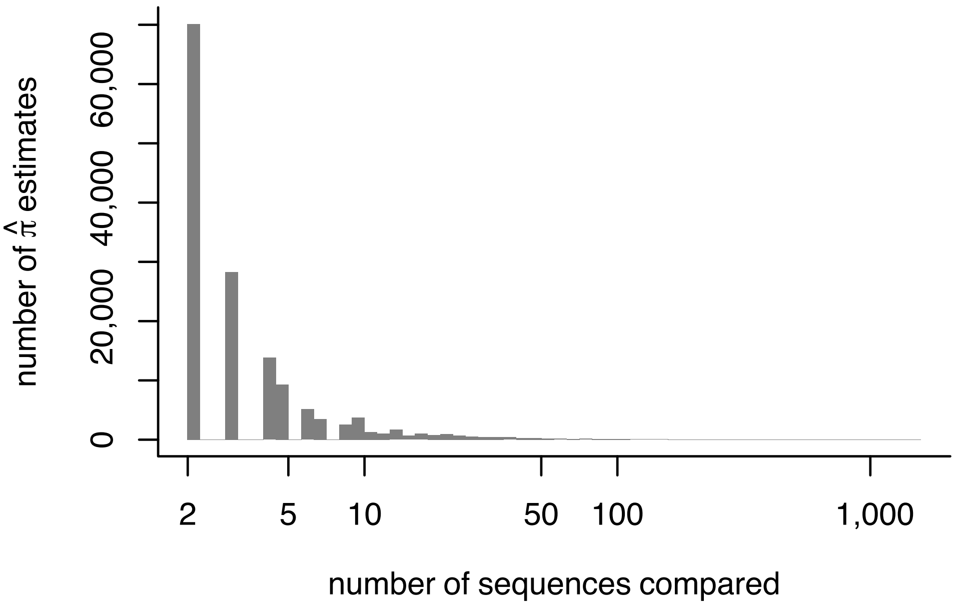

*Figure S7*. Results of sensitivity analysis manipulating the minimum number of sequences required to compute a diversity estimate. This figure is analogous to Fig. 3 of the main text, but in this case for GAMMs only using diversity estimates computed from > 5 sequences. Figure design is identical to Fig. 3 of the main text.

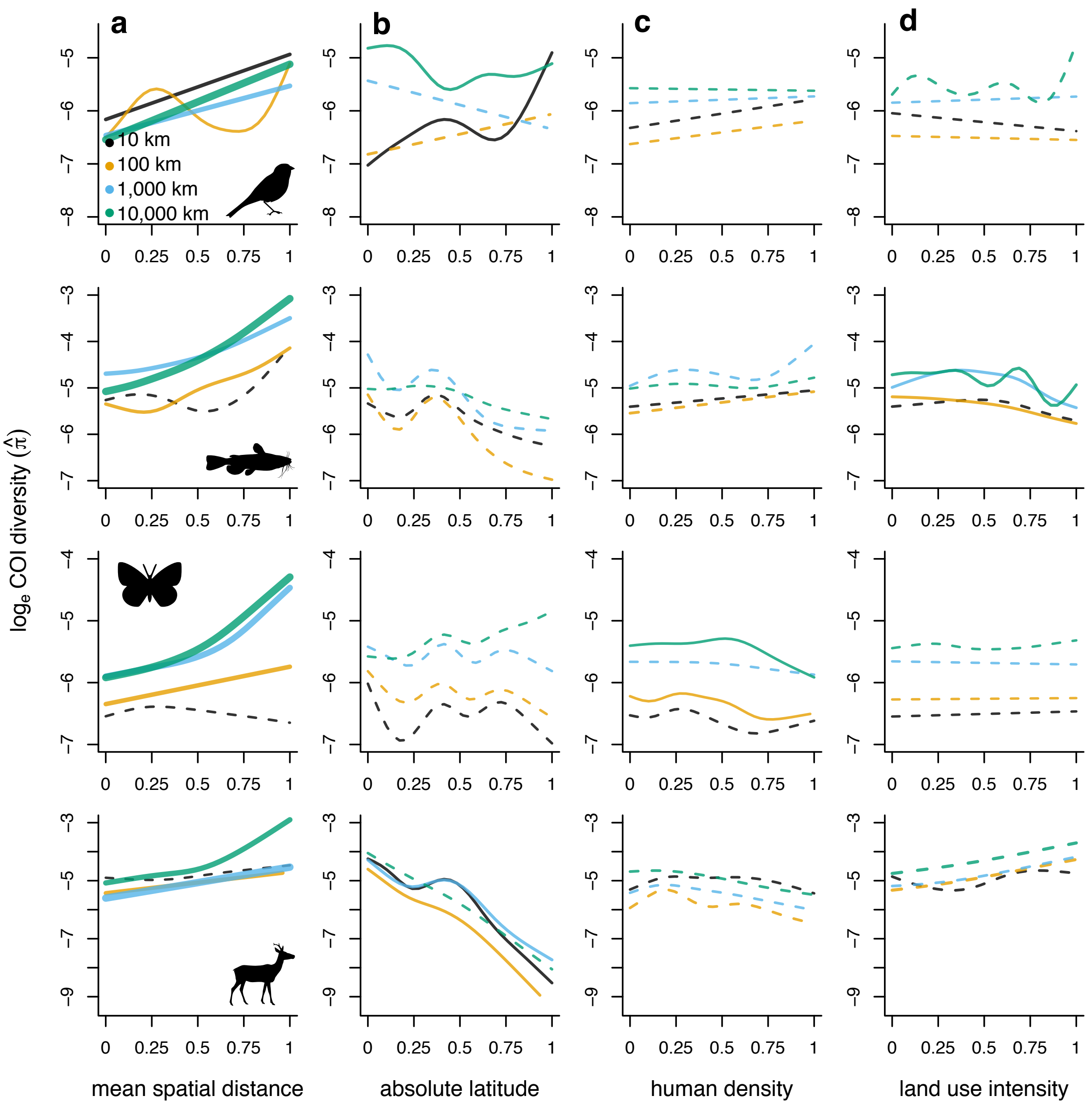

*Figure S8.* Results of spatial GAMMs without weights but including as a fixed effect/smooth term the log-transformed number of sequences on which each diversity estimate is based. Figure design is identical to Fig. 3 of the main text.

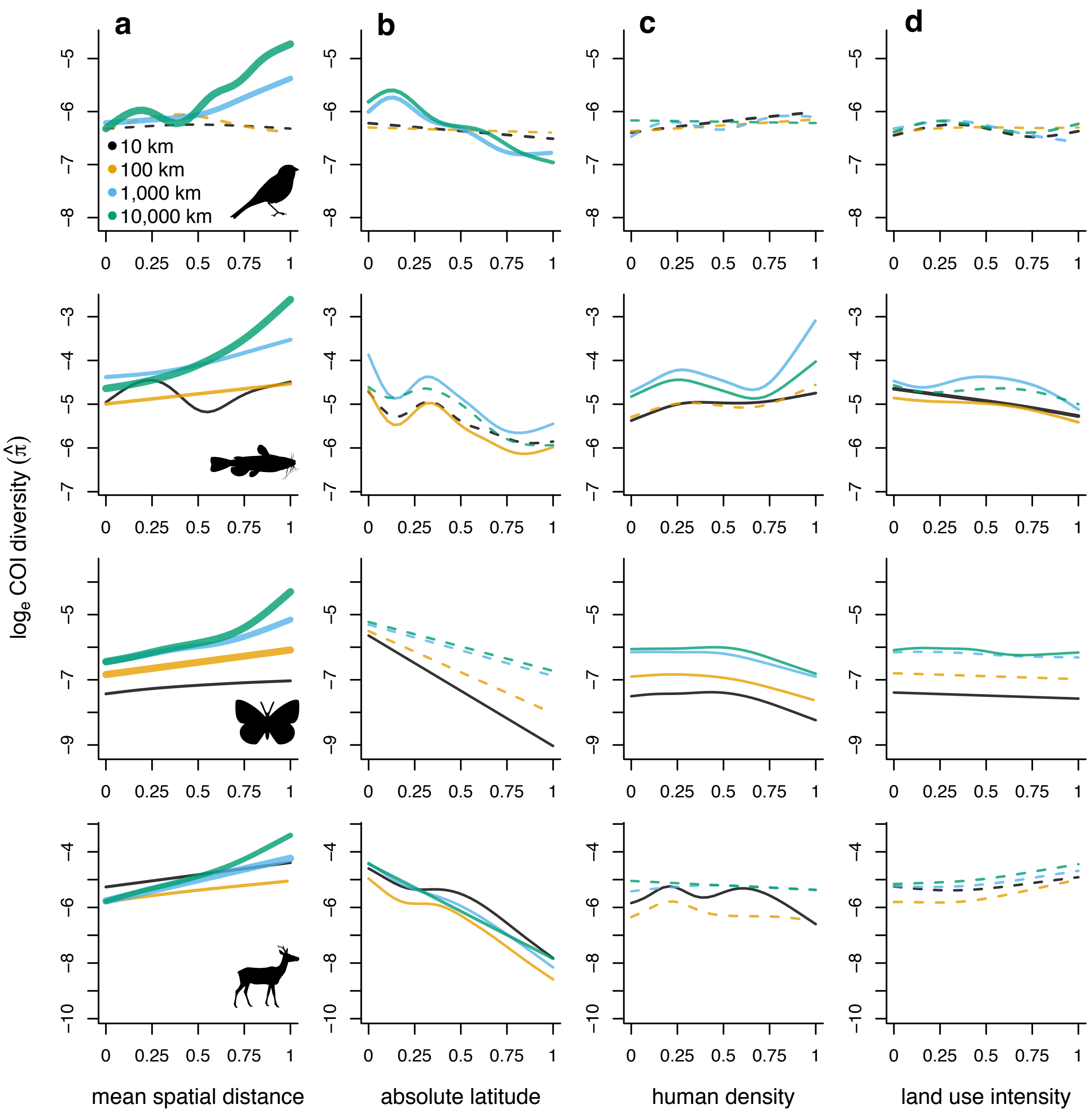

*Figure S9.* Results of spatial GAMMs including 'hemisphere' as a parametric effect and an 'hemisphere × latitude' factor-smooth interaction. Predicted values for both hemispheres were obtained while setting values for other predictors to their median value in the scale and taxon-specific dataset. The full structure of the model is provided in Table S1. Panels from the same column belong to the same taxon. For each model, the *p* value of the 'difference smooth' is provided to indicate whether trends are significantly-different between the two hemispheres.

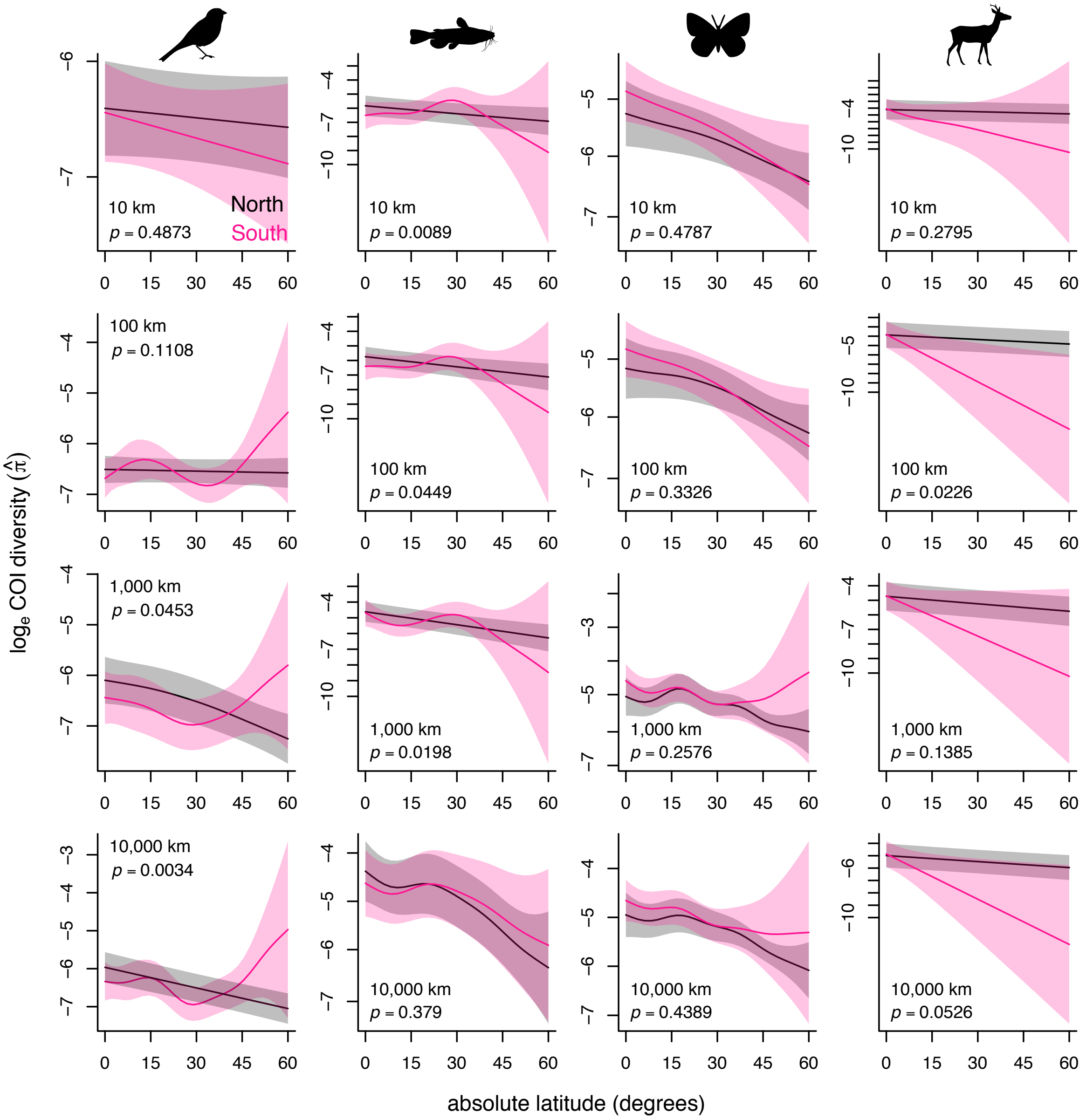

*Figure S10.* Effects of human population density on genetic diversity in spatial GAMMs including 'biome' as a parametric effect and a 'biome × latitude' factor-smooth interaction. Predicted values for both biomes were obtained while setting values for other predictors to their median value in the scale and taxon-specific dataset. The full structure of the model is provided in Table S1. Panels from the same column belong to the same taxon. For each model, the *p* value of the 'difference smooth' is provided to indicate whether trends are significantly-different between the two biomes.

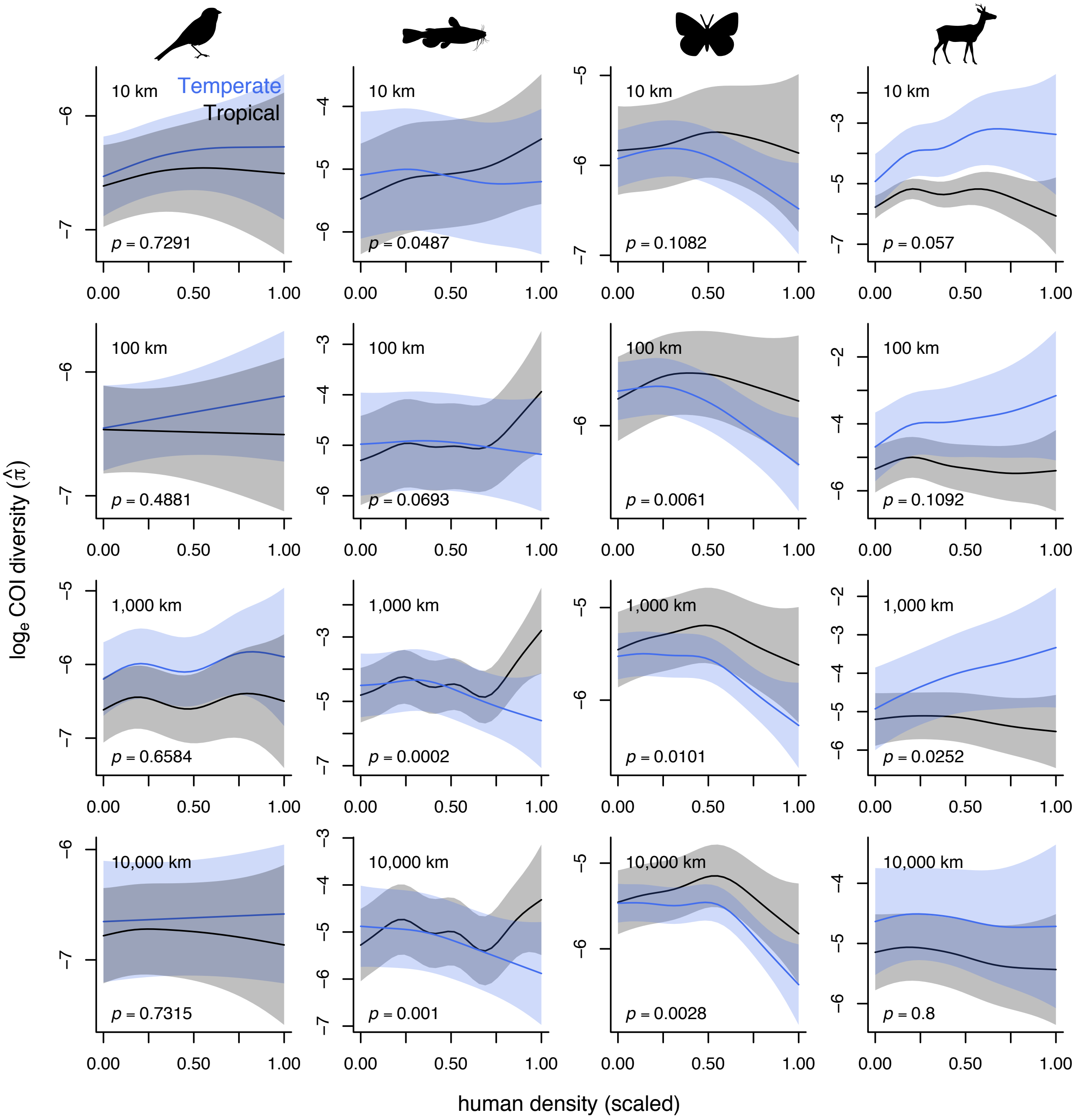

*Figure S11.* Effects of land use intensity on genetic diversity in spatial GAMMs including 'biome' as a parametric effect and a 'biome × latitude' factor-smooth interaction. Predicted values for both biomes were obtained while setting values for other predictors to their median value in the scale and taxon-specific dataset. The full structure of the model is provided in Table S1. Panels from the same column belong to the same taxon. For each model, the *p* value of the 'difference smooth' is provided to indicate whether trends are significantly-different between the two biomes.

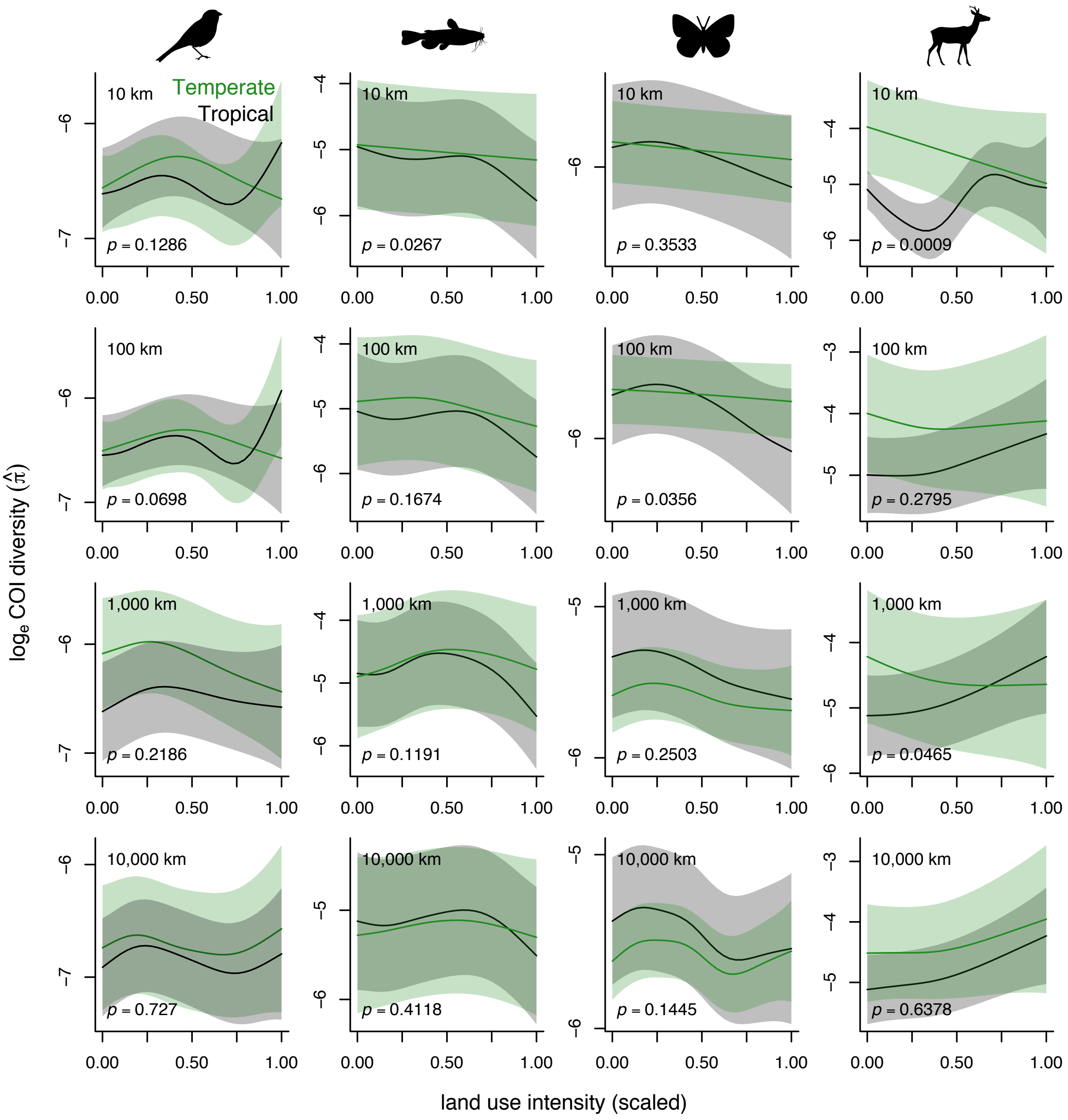

*Figure S12.* Results of time series GAMMs without weights but including as a fixed effect/smooth term the log-transformed number of sequences on which each diversity estimate is based. Figure design is identical to Fig. 5 of the main text.

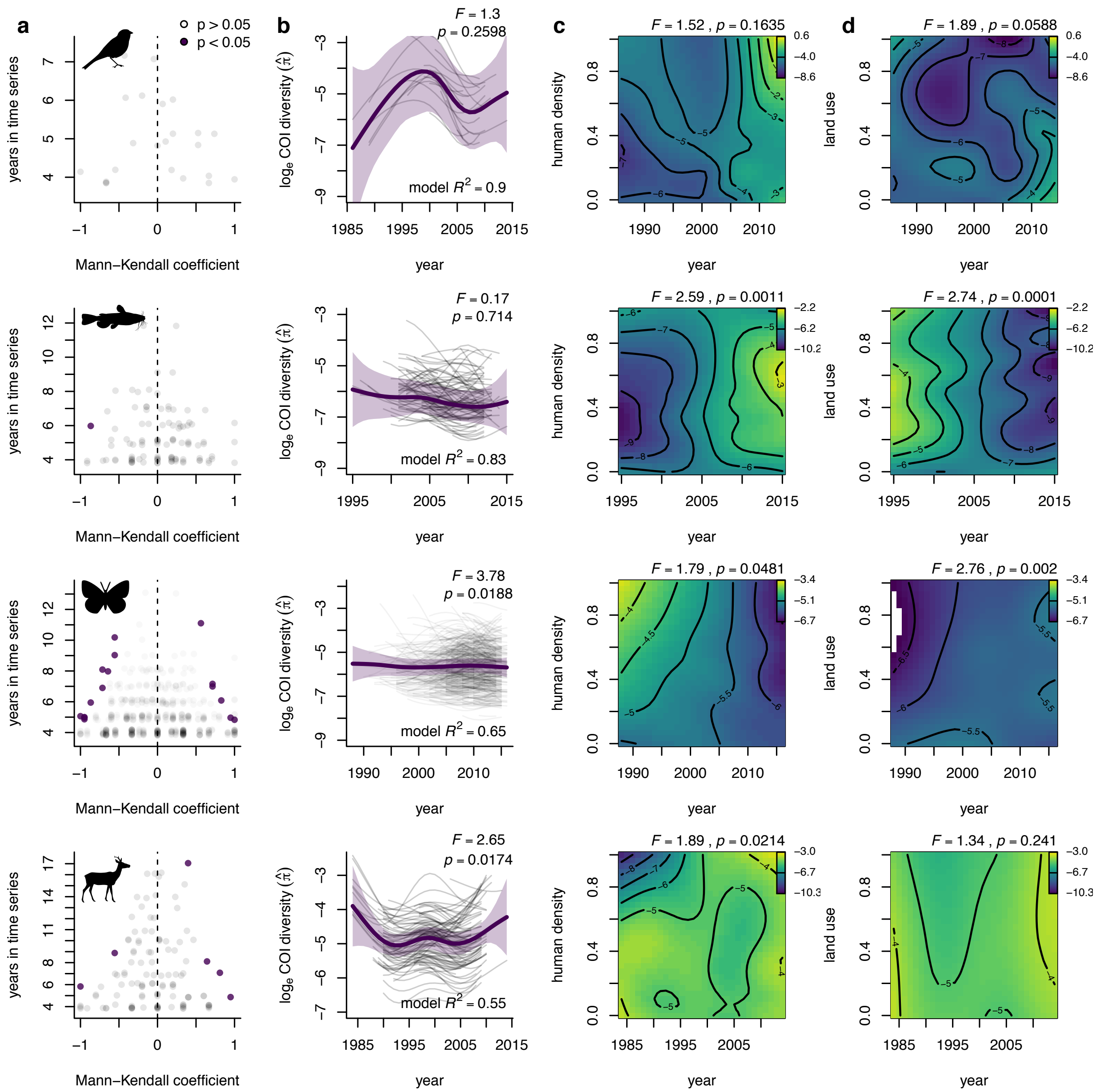
